# A FeuP–FeuQ–Dependent Regulatory Module Integrating Envelope Stress and Host-Associated Behaviors in *Agrobacterium tumefaciens*

**DOI:** 10.64898/2026.01.20.700542

**Authors:** Michelle C. Brock, Daniel Arango-cardona, Jordan N. Pantelione, Jason E. Heindl

**Author notes:** Address correspondence to Jason E. Heindl,.

## Abstract

*Agrobacterium tumefaciens* is an important plant pathogen, the causative agent of crown gall disease, and a foundational technology used for genetic transformation of plant tissue. More recently, *A. tumefaciens* has been adopted as a genetically tractable model organism for studying bacterial cell cycle regulation, developmental pathways, and niche construction. The transition from a free-living bacterium within the rhizosphere to association with a plant host and subsequent transformation of plant tissue is one aspect of *A. tumefaciens*’ life history that encompasses all three of these processes. Such events are coordinated by multiple regulatory modules which must sense external and/or internal cues, integrate these inputs, and effect appropriate changes in gene expression and cellular responses to maximize fitness. Here, we evaluate the contribution of the two-component system, FeuP-FeuQ, to gene expression and developmental phenotypes including surface attachment/biofilm formation, swimming motility, and tumorigenesis. *feuPQ* operon organization suggests translational coupling during expression of the response regulator, FeuP (Atu0970), and sensor kinase, FeuQ (Atu0971). In-frame, non-polar deletion of *feuP* or *feuQ* individually, or the entire *feuPQ* operon, resulted in reduced biofilm formation, swimming motility, and tumor formation, without adversely affecting planktonic growth. Transcriptomic profiling identified ∼300 differentially expressed genes when the *feuPQ* locus was disrupted, including genes affecting flagellar motility, succinoglycan production, and type VI secretion. Phenotype profiling emphasized the contribution of *feuPQ* to withstanding osmotic, ionic, and antimicrobial stressors. Together, these data highlight FeuPQ as a global regulator of cellular responses which likely contribute to overall fitness during rhizosphere lifestyle transitions.

**IMPORTANCE:** *Agrobacterium tumefaciens* is an important plant pathogen able to genetically transform numerous plant species. In the rhizosphere, this bacterium encounters many challenges ranging from antagonistic and competing microbes, to plant host defenses, to rapidly changing environmental conditions. Efficient host interaction leading to plant transformation requires coordination of bacterial motility, attachment, and defense mechanisms, among other processes. This work identifies two proteins, FeuP and FeuQ, that together contribute to such coordination. The importance of this work is in identifying the FeuPQ system as a global regulator of these and other processes, possibly enabling targeted interventions to promote or inhibit plant transformation.

## INTRODUCTION

Adaptation to changing environments requires coordinated regulatory programs that enable bacteria to sense, integrate, and respond to diverse physiochemical cues. Environmental signals such as pH, osmolarity, nutrient limitations, or plant-derived signals are continuously monitored by bacteria to enable rapid changes. Survival tactics required by the constant changes are essential for bacterial survival across niches, especially for organisms that transition between different habitats (1, 2).

*Agrobacterium tumefaciens (A. tumefaciens*), a soil dwelling member of the *Alphaproteobacteria,* is a well-established model for studies of bacterial signaling networks and resultant phenotypes. This is due to its ability to transition between two distinct lifestyles: a sessile, surface-associated state and a free-swimming, motile state. In the sessile state, *A. tumefaciens* cells attach to biotic or abiotic surfaces, often as part of a structured multicellular biofilm. Biofilm formation enhances the chances of *A. tumefaciens* to display increased resistance to stress and increases opportunity for metabolic cooperation (3, 5). On the contrary while in its motile state, *A. tumefaciens* are actively navigating through different environments. The switch between sessile to motile state is tightly regulated, indicating that it’s crucial for successful colonization (3, 4, 6).

Central to regulatory control are two-component regulatory systems (TCS). TCS are modular signaling systems that generally consist of two conserved proteins: a membrane associated histidine kinase and a response regulator. The canonical histidine kinase (HK) is a protein that will detect environmental stimuli, to initiate signal transduction. The response regulator (RR) serves as the output signal, frequently altering gene expression. Upon sensing a specific input, the HK undergoes autophosphorylation at a conserved histidine residue. The resulting phosphate group is transferred to an aspartic acid residue, located within the REC domain of the RR. Although TCS are commonly described as linear, one-to-one signaling pathways, variations such as hybrid histidine kinases, multi-step phosphorelays and cross-regulation among multiple systems are common, particularly in complex bacterial genomes (1, 2, 7–13).

In *A. tumefaciens*, TCS are crucial for regulating a wide array of adaptive behaviors. These behaviors include: biofilm formation, motility, cell cycle progression, and virulence. To date, at least 25 two-component systems have been identified (16, 20) in *A. tumefaciens*. Genome annotation suggests that *A. tumefaciens* encodes dozens of histidine kinases and response regulators, highlighting the complexity of its two-component signaling capacity. These TCS systems are distributed across multiple replicons in *A. tumefaciens*. This likely contributes to coordinating signals that govern both environmental adaptation and host-pathogen interactions.

Among the best characterized systems is VirA-VirG. This system is located on the Ti plasmid and senses plant-derived phenolic compounds such as acetosyringone and activates expression of *vir* genes. The expression of *vir* genes are necessary for T-DNA transfer and transformation of plant host cells (6, 12, 13). Another well-known two-component regulatory system is the ChvG-ChvI system. This system, which is encoded chromosomally, plays a key role in the acid stress response. It also participates in the regulation of host colonization, including virulence factor production and exopolysaccharide synthesis. Under acidic or envelope stress conditions, ChvG is activated, leading to phosphorylation of ChvI. Once ChvI is phosphorylated, transcriptional induction of genes promoting bacterial survival or tumorigenesis occurs (14, 16). A more complex system incorporating two-component signaling, the PdhS-DivK-CtrA regulatory pathway, is a phosphorelay involved in the regulation of cellular differentiation and development. This system plays a role in pole differentiation, motility, and DNA replication. Disruption of these TCS result in profound alterations in bacterial morphology and the overall fitness of *A. tumefaciens* (3, 7–9, 11, 14, 17–18).

Despite significant advances in our understanding of key TCS pathways, many predicted TCS in *A. tumefaciens* remain functionally uncharacterized. One incompletely studied two-component regulatory locus is *feuP-feuQ*. The *feuP-feuQ* operon is conserved across members of the *Rhizobiales* order, yet its functional role in *A. tumefaciens* has not been investigated (41). The *feuP-feuQ* locus encodes two putative signaling proteins whose domain organization is consistent with a canonical TCS. Senor histidine kinase, FeuQ, is predicted to carry two transmembrane helices, a cytoplasmic HisKA domain responsible for autophosphorylation and a HATPase_c domain that mediates ATP binding and phosphate transfer. Sequence alignment suggests autophosphorylation activity occurs at a conserved histidine residue at position 255. Its presumed cognate response regulator, FeuP, harbors a receiver (REC) domain with a conserved aspartic acid residue at position 51, and a carboxy-terminal OmpR/PhoB-type DNA-binding output domain, assumed to function in transcriptional regulation. Phosphorylation of this conserved aspartic acid is predicted to promote DNA binding and transcriptional regulation by FeuP. Based on these features, we hypothesize that FeuP-FeuQ functions as a classic two-component regulatory system that senses one or more environmental signals, thus enabling adaptation of the bacterium and regulating phenotypes essential for survival and host interaction (7, 9, 24).

In this study, we investigate the biological role of FeuP-FeuQ on the regulation of a set of fundamental developmental phenotypes in *A. tumefaciens*. To dissect the regulatory function of this system, we used a combination of genetic, phenotypic, and molecular approaches. Deletion and overexpression strains were constructed to evaluate the impact of FeuP-FeuQ on bacterial swimming motility, biofilm formation, virulence, and growth. RNA sequencing (RNA-seq) and phenotypic microarray analysis were used to identify differentially expressed genes and growth phenotypes, respectively, in Δ*feuP*, Δ*feuQ*, and Δ*feuPQ* backgrounds. Together these experiments reveal that FeuP-FeuQ constitutes a previously uncharacterized signaling pathway involved in coordinating *A. tumefaciens* developmental phenotypes in response to environmental signals (5, 17, 19).

## RESULTS

### Characteristics of the *feuP-feuQ* locus, encoding a canonical two-component regulatory system

The *feuP-feuQ* locus includes an overlapping set of two genes under control of the same promoter, with *feuP* (*atu0970*) immediately upstream of *feuQ* (*atu0971*). The primary transcription start site is located 57 base pairs upstream of the *feuP* start codon (Fig.1A). The last 11 base pairs of the *feuP* coding sequence overlap with the beginning of the *feuQ* coding sequence, suggesting translational coupling of the gene pair encoded on this operon. Immediately downstream of the *feuP-feuQ* operon is a predicted surface protein encoded by *lipA* (*atu0972*), transcribed independently from its own promoter. Immediately upstream is the coding sequence for a small hypothetical protein (Atu8142), also transcribed from its own promoter (42). Domain architecture predictions are consistent with a classical histidine kinase-response regulator pair.

**Fig 1.**
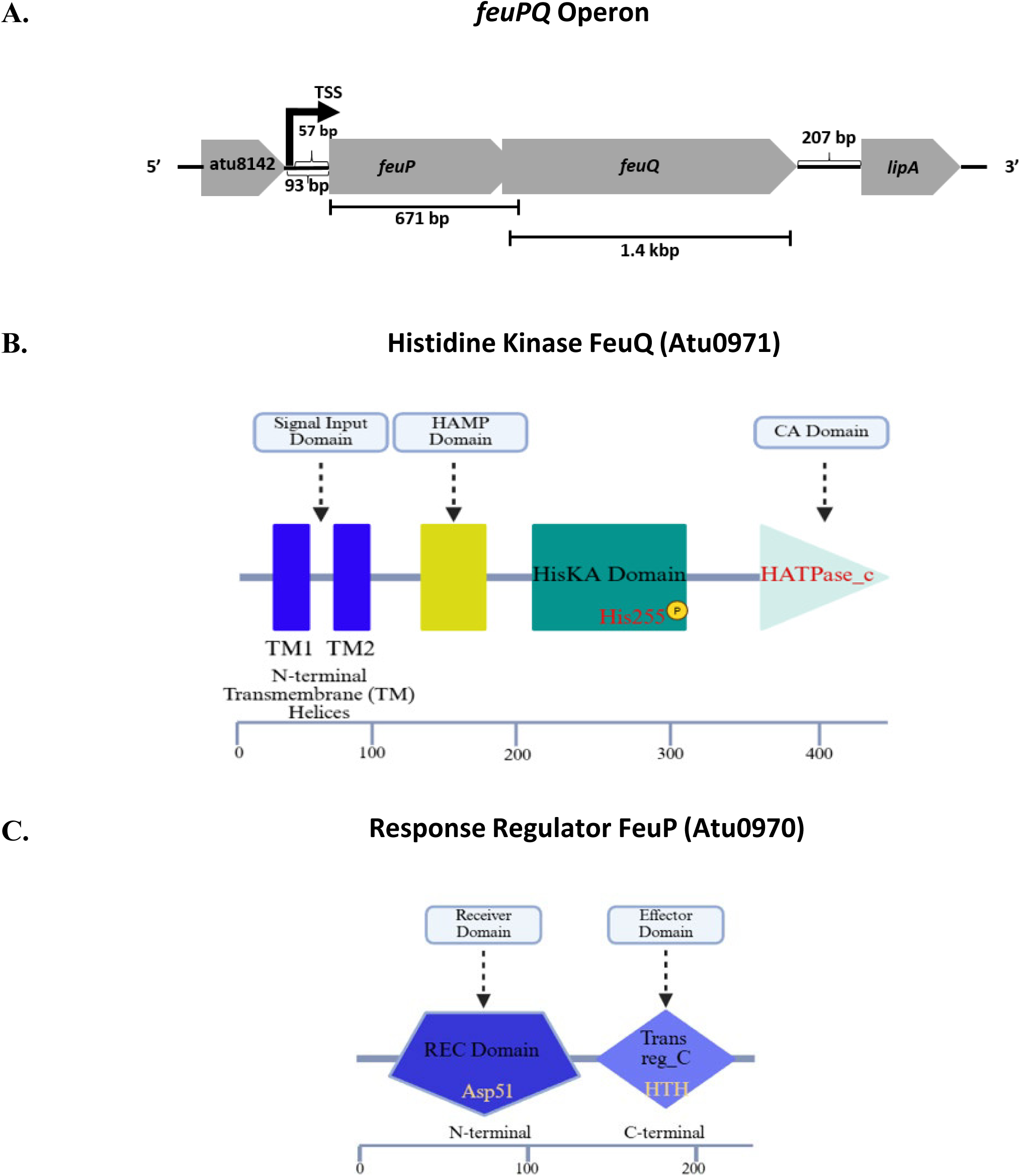
Genomic and protein domain organization of FeuP-FeuQ two-component regulatory system. **(A**) Genomic organization of the *feuP-feuQ* (*atu0970*-*atu0971*) operon. Arrow indicates transcriptional start site from (42) . There is an overlap of 11 base pairs for the coding sequences of *feuP* and *feuQ*. *atu8142* is located 93 base pairs upstream. *lipA* (*atu0972*), encoding a predicted surface protein, LipA, is 207 base pairs downstream. **(B)** Predicted domain architecture of FeuQ (inner membrane histidine kinase). FeuQ contains two transmembrane helices at the amino terminus (blue bars), a cytosolic HisKA domain, and a HATPase_c domain at the carboxyl terminus. **(C)** Predicted domain architecture of the response regulator FeuP. FeuP contains an amino-terminal receiver (REC) domain and a carboxy-terminal helix-turn-helix (HTH) DNA binding domain. Models were adapted from SMART Database (http://smart.embl.de). Numbering beneath protein domain models in B and C indicate amino acid length. Biorender (https://app.Biorender.com) was used to generate components of the illustration.

FeuQ is predicted to be an inner membrane histidine kinase with two N-terminal transmembrane helices and cytosolic HisKA and HATPase_c domains (Fig. 1B). The HisKA region includes the conserved eponymous histidine residue onto which autophosphorylation occurs, and the HATPase_c contains conserved N, G1, F, G2 nucleotide-binding motifs, which support ATP-dependent phosphotransfer chemistry (43,44). FeuP is predicted to be a cytosolic response regulator that contains an N-terminal receiver (REC) domain and a C-terminal helix-turn-helix DNA binding domain (Fig. 1C). The REC domain retains the established acidic residues surrounding the phospho-acceptor aspartate and the catalytic Lys/Thr pair, typically seen in the OmpR/PhoB like regulators. This is consistent with phosphorylation dependent activation of the downstream DNA binding domain (13).

### FeuP-FeuQ signaling does not determine growth under standard conditions

To study the role of the FeuP-FeuQ two-component system in *A. tumefaciens*, initial characterization began with growth and the gross developmental phenotypes of surface attachment and motility, considered standard phenotypes used by the bacterium for persisting in the environment. The *feuP* and *feuQ* coding sequences were removed from the genome, singly and in combination, using a plasmid-based homologous recombination protocol. All resultant strains were viable and displayed no obvious colony phenotype on ATGN plates. To explore growth kinetics, liquid cultures of the wild-type strain C58 and its isogenic deletion strains, Δ*feuPQ*, Δ*feuP*, and Δ*feuQ*, were monitored by optical density (Fig. S1). Across the full course of time totaling 36 hours, growth overlapped with comparable lag durations and similar doubling time during the exponential phase. We also observed indistinguishable time to stationary phase between the four constructs. Statistical analysis did not detect strain-dependent differences in the final turbidity at any time point. We conclude that neither *feuP* nor *feuQ* are essential for growth on ATGN, and that disruption of the *feuP-feuQ* locus does not produce a detectable proliferation defect. These results accord with prior work that did not identify either *feuP* or *feuQ* as essential for growth (45).

### FeuP-FeuQ promotes efficient surface attachment

To evaluate the role of FeuP-FeuQ in surface attachment, we performed static biofilm assays using plasmid-borne alleles of *feuP* and *feuQ* for ectopic expression, and the deletion strains referenced above. Overexpression of *feuQ*, but not *feuP*, from an IPTG-inducible promoter resulted in a significant increase in biofilm formation in the wild-type background. Loss of either *feuP* or *feuQ* resulted in a significantly reduced attachment relative to the wild-type strain (Fig. 2A). Plasmid-based expression of *feuQ* significantly increased biofilm formation in the Δ*feuQ* background, but did not complement the phenotype to wild-type levels. Plasmid-based expression of *feuP* did not significantly increase biofilm formation in the Δ*feuP* background. Based on these results, the Δ*feuPQ* double mutant was evaluated, with the hypothesis that for efficient activity of the FeuP-FeuQ system the two genes must be co-expressed. Simultaneous loss of both *feuP* and *feuQ* in the Δ*feuPQ* double mutant reduced biofilm formation to levels similar to either the Δ*feuP* or Δ*feuQ* strains. Plasmid-based expression of either *feuP* or *feuQ* in the Δ*feuPQ* double mutant failed to restore biofilm formation in this background. Plasmid-based expression of the complete *feuPQ* operon, however, restored biofilm levels, even providing an enhanced accumulation of adherent biomass compared to the wild-type strain. An increase in biofilm formation was also evident when the *feuPQ* operon was overexpressed in the wild-type strain background. Collectively, these results support a positive role for FeuP-FeuQ in surface attachment under the conditions assayed, and suggest that expression of *feuP* is tightly coupled with that of *feuQ*.

**Fig 2.**
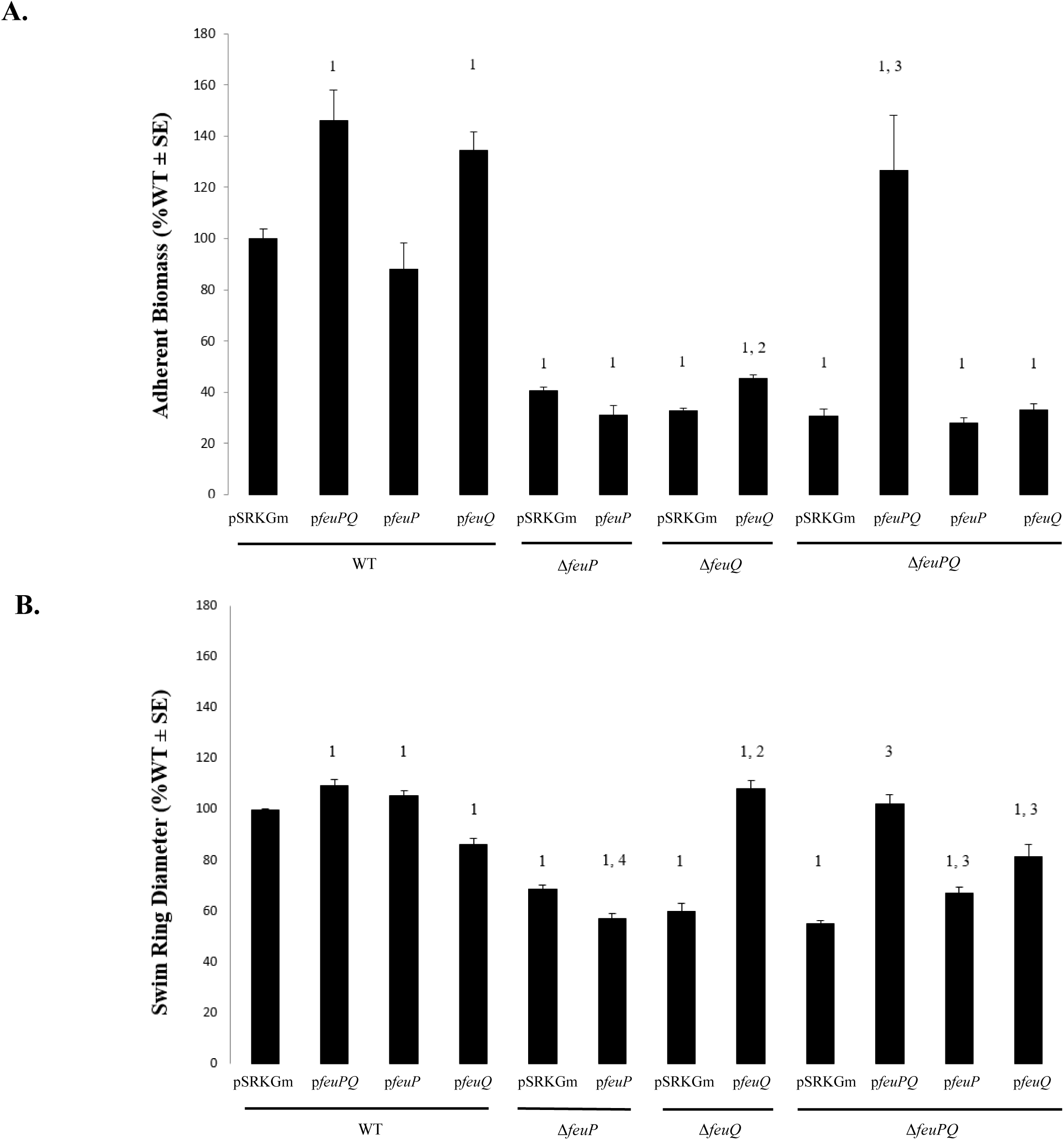
**Loss of *feuP* or *feuQ*, singly or in combination, significantly alters surface attachment and motility phenotypes**. **(A)** Static biofilm formation assays were performed to assess attachment to plastic coverslips over the span of 48 hours of the indicated strains carrying empty vector, pSRKGm, of plasmid-borne wild-type alleles of *feuP*, *feuQ*, or the entire *feuPQ* operon. Adherent biomass was calculated as the ratio of biofilm biomass to planktonic growth of cultures. Biofilm biomass was determined by solubilizing biofilm-adsorbed crystal violet and measuring the absorbance at 600 nm. Planktonic growth of biofilm cultures was measured as optical density (OD_600_). All strains were grown in ATGN media, induced with 100 μM IPTG. **(B)** Plate-based swim motility assays were performed to assess generic motility for a total period of 7 days. Strains were inoculated in 0.25-0.30% agar ATGN plates, induced with 100 μM IPTG. Swim ring diameters were measured daily. Values for both assays were normalized to the wild-type strain bearing an empty vector. Bars show mean ± SE from three independent biological replicates (n = 9). Statistics were calculated using Welch’s T-Test: Two-Tailed, Unequal Variance, compared to the wild-type strain bearing an empty vector. ^1^P < 0.05 compared to wild-type C58 pSRKGm, ^2^P < 0.05 compared to Δ*feuQ* pSRKGm, ^3^P < 0.05 compared to Δ*feuPQ* pSRKGm, ^4^P < 0.05 compared to Δ*feuP* pSRKGm

### FeuP-FeuQ supports efficient swimming motility

A plate-based motility assay was used to evaluate any contribution of the FeuP-FeuQ system to motility. Overexpression of either *feuP*, *feuQ*, or *feuPQ* from an IPTG-inducible promoter resulted in modest but statistically significant changes in swim ring diameter in the wild-type background. Both the Δ*feuP* and Δ*feuQ* strains were significantly reduced in their ability for colonies to expand outwardly from the inoculation site, as measured by diameter of swim rings after seven days (Fig. 2B). As seen for surface attachment, motility was efficiently complemented by plasmid-based expression of *feuQ* of the Δ*feuQ* background. Also as seen with surface attachment, plasmid-based expression of *feuP* failed to complement the motility defect of the Δ*feuP* background. The swim ring diameter produced by the Δ*feuPQ* double mutant strain was significantly reduced compared to the wild-type strain, and of a similar reduction compared to either the Δ*feuP* or the Δ*feuQ* single mutants. This reduction in swim ring diameter was efficiently complemented by plasmid-based expression of the intact *feuPQ* operon. In contrast to surface attachment, plasmid-based expression of either *feuP* or *feuQ*, singly, significantly increased swim ring diameters in this background, although not with the same efficiency as the complete *feuPQ* operon. These results suggest that motility as measured using this plate-based assay is regulated by the FeuP-FeuQ system.

### FeuP-FeuQ is required for efficient tumor formation on plant tissue

Surface attachment and motility are two broad functions contributing to *A. tumefaciens* pathogenesis. As an initial step in determining if, and to what degree, the FeuP-FeuQ system influences host interaction, plant transformation, and plant tumor formation, we employed a potato disk assay. Equivalent culture densities of the wild-type, single mutant (Δ*feuP* or Δ*feuQ*), and double mutant (Δ*feuPQ*) strains, were individually inoculated onto sterile potato disks and monitored for tumor formation over the course of three weeks. All strains lacking either *feuP* or *feuQ* exhibited a significant reduction in the total number of tumors formed relative to the wild-type strain (Fig. 3). The Δ*feuPQ* strain showed the most pronounced reduction, while the single deletions yielded comparable phenotypes in reduction of tumor number. These results suggest that FeuP and FeuQ act together to support full virulence.

**Fig 3.**
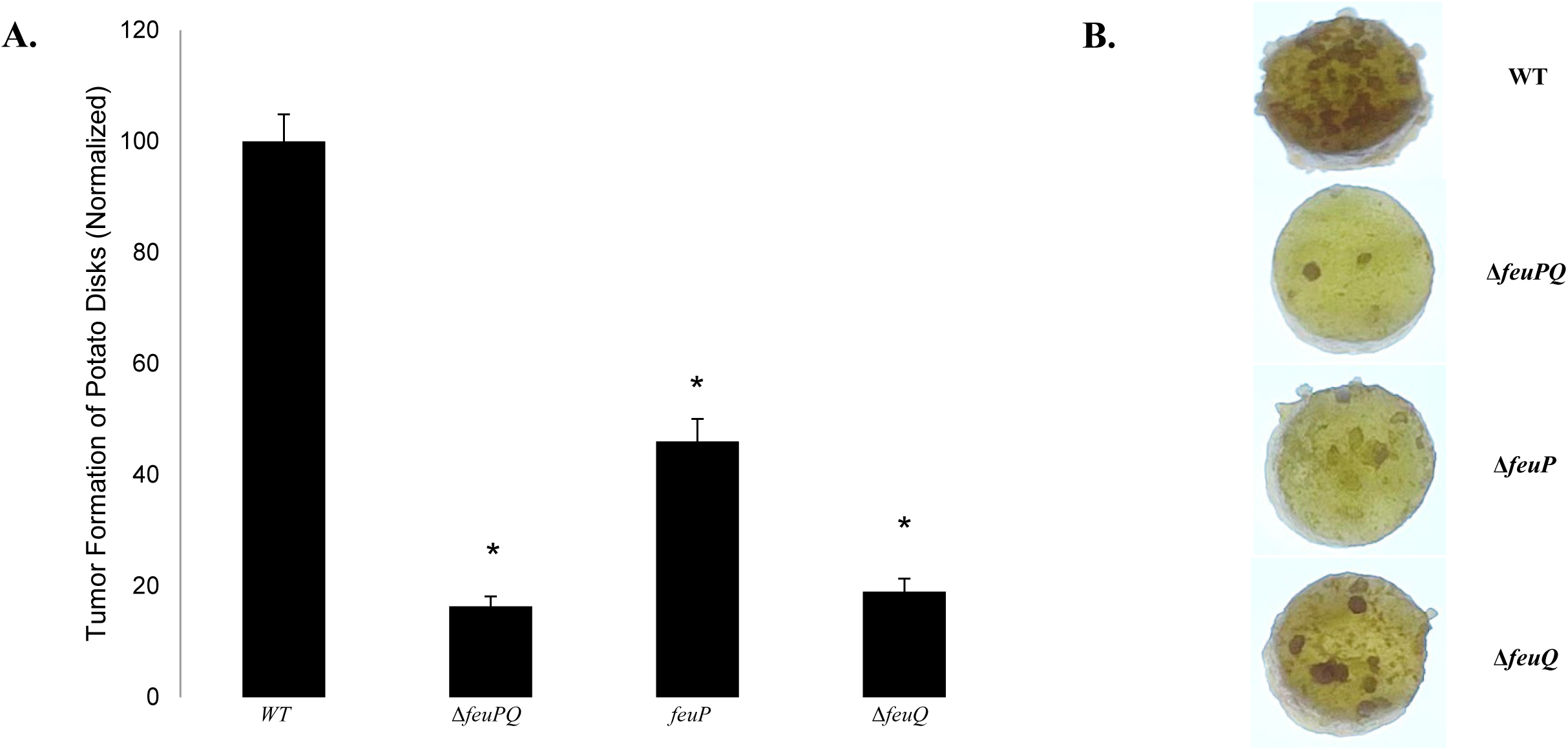
***feuP* and *feuQ* are required for robust tumor formation.** Sterile potato discs were inoculated with the indicated strains and incubated for a total period of 21 days. Tumors formed post inoculation were counted on day 21. **(A)** Quantification of tumor counts, normalized to the mean number of tumors formed by the wild-type strain (%WT). Bars show mean ± SE from three independent biological replicates (n = 9). All mutants exhibited significantly reduced tumor formation relative to the wild-type strain. Statistics were calculated using Welch’s T-Test: Two-Tailed, Unequal Variance. Asterisks denote p values where p < 0.0001. **(B)** Representative images of potato discs at day 21 post inoculation for the indicated strains. Discs were imaged under identical settings. Brightness and contrast were adjusted uniformly across all panels.

### Transcriptome profiling implicates FeuP-FeuQ as a global regulator

To characterize the transcriptional effects of disrupting the FeuP-FeuQ two-component regulatory system, we performed transcriptome profiling using RNA sequencing (RNA-seq) on the wild-type strain and the three isogenic deletion strains, Δ*feuPQ,* Δ*feuP* and Δ*feuQ.* Differentially expressed genes (DEGs) were defined in comparison to wild-type levels of gene expression under the experimental conditions. For analysis, genes exhibiting either a two-fold or greater increase or decrease in transcript abundance were binned as DEGs. Loss of either *feuP* or *feuQ*, alone or in combination, resulted in large changes in gene expression. Loss of *feuP* resulted in 165 DEGs, 71 of which were upregulated and 94 downregulated. Loss of *feuQ* resulted in 133 DEGs, with expression of 46 genes upregulated and 87 genes downregulated. Absence of the complete *feuPQ* operon resulted in 188 DEGs, 62 of which were upregulated and 126 of which were downregulated. Volcano plots of the DEGs for each strain show a wide range of magnitude of expression changes for all three strain backgrounds (Fig. 4A, S2).

**Fig 4.**
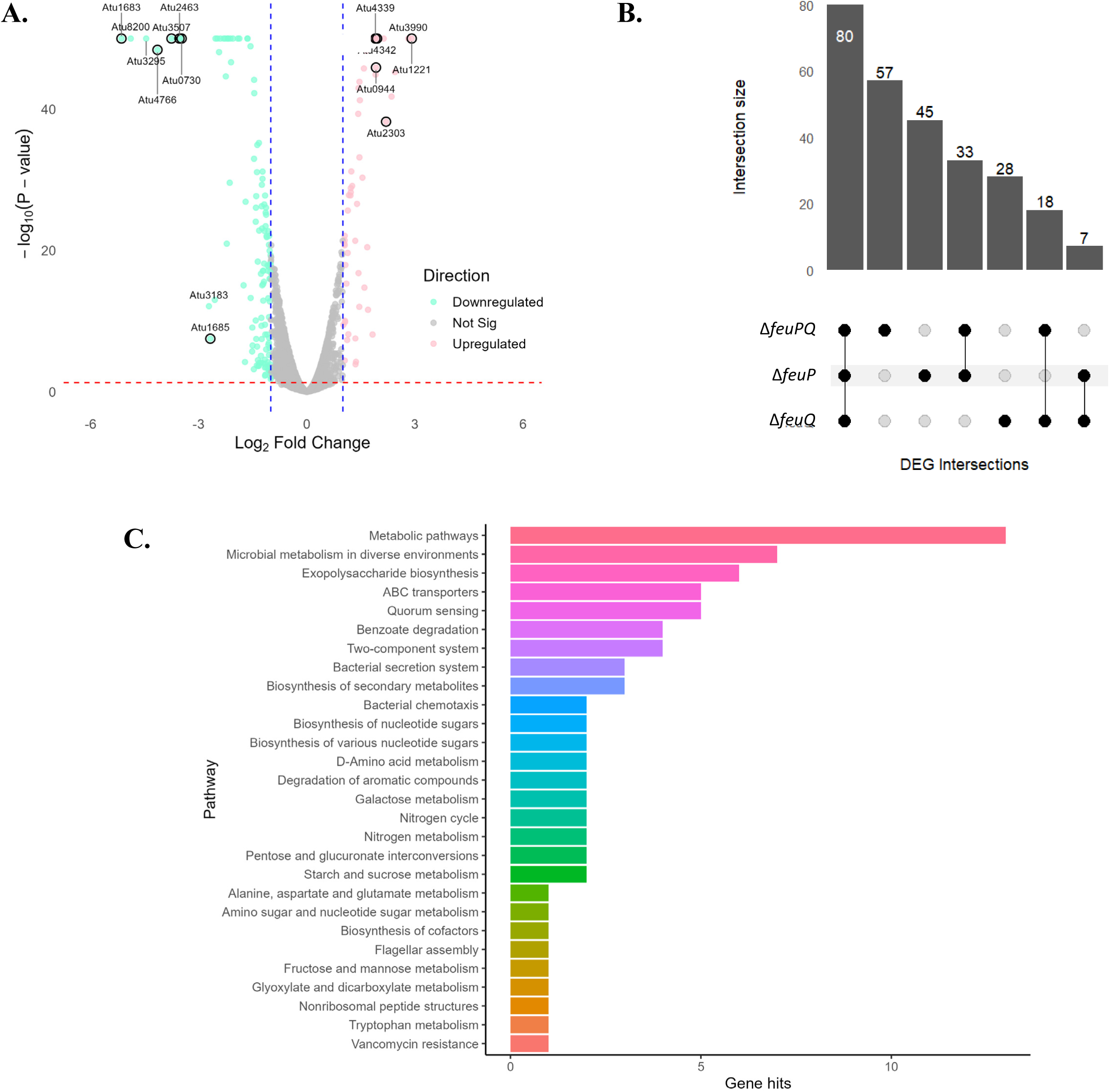
Loss of *feuP* or *feuQ* elicits overlapping and mutant-specific transcriptional changes. **(A)** Volcano plot for the suite of genes which are differentially expressed in the Δ*feuPQ* background, relative to the wild-type background. RNA-seq was performed on three biological replicates. Blue vertical lines indicate a log_2_ (fold-change) of ± 1. Red horizontal line represents a p value < 0.05. A total of 188 genes were differentially expressed, with 126 downregulated (green dots) and 62 upregulated (pink dots). **(B)** Complex upset plot demonstrating the intersection of the suites of differentially expressed genes (DEGs) identified for the indicated strains, relative to wild-type gene expression. Volcano plots for the Δ*feuP* and Δ*feuQ* DEGs are included in the supplemental material. **(C)** Bar plots display the number of DEGs assigned to each Kyoto Encyclopedia of Genes and Genomes (KEGG) pathway, ranked by gene count for the Δ*feuPQ* background. KEGG pathway lists for the Δ*feuP* and Δ*feuQ* DEGs are included in the supplemental material.

There is considerable overlap between the three DEG sets (Fig. 4B, Tables 1-3, S4–S6). Of 268 total DEGs identified among all three strain backgrounds, 80 (30%) are differentially expressed in all three strain backgrounds. An additional 58 (22%) are shared between two of the strain backgrounds. The overlap in DEGs is demonstrated when comparing the set of twenty loci which are most highly differentially expressed in each background, selecting the top 10 upregulated and top 10 downregulated genes for each (Tables 1-3). In this case, only four genes appear as uniquely regulated in only one background (*atu3183*, *copA*, *copB*, and *traC*). Among the overlapping DEGs in this restricted dataset are a number of genes involved in type VI secretion (*impB*, *impC*, *impE*, *impK*, *impJ*, *impL*), and potential surface-associated structures such as curli (*atu4766*, *atu4767*, *atu4768*) and lectins (*atu0730*, *atu3295*). In all three strains, the most highly upregulated gene was *atu1221*, encoding a hypothetical protein belonging to the NlpC/p60 family, containing a signal peptide and a putative endopeptidase domain. Taken together, these data demonstrate that FeuP and FeuQ influence a broad transcriptional program in *A. tumefaciens* with shared and potentially independent regulatory roles (28).

**Table 1.** Summary of top genes upregulated and downregulated in the Δ*feuPQ* mutant identified by transcriptome analysis. RNA-seq comparison between *A. tumefaciens* wild-type and the Δ*feuPQ* mutant revealed genes with an increase or decrease in transcript abundance.

| Locus | Gene Name | Predicted Function | logFC |
| --- | --- | --- | --- |
| Atu1221 |  | NlpC/p60 superfamily protein | 2.913717 |
| Atu3990 | *copC* | Copper Tolerance protein | 2.817807 |
| Atu3992 | *copB* | Copper Tolerance protein | 2.448915 |
| Atu3991 | *copA* | Multicopper oxidase | 2.354589 |
| Atu2303 |  | Polysaccharide deacetylase | 2.201484 |
| Atu1805 |  | BON domain-containing protein | 2.136338 |
| Atu4342 | *impB* | Type VI secretion | 1.952352 |
| Atu0944 | *cscA* | Sucrose hydrolase | 1.922206 |
| Atu0898 |  | DUF930 domain-containing protein | 1.910969 |
| Atu4339 | *impE* | Type VI secretion | 1.906814 |
| Atu1835 |  |  | -2.55173 |
| Atu1685 |  |  | -2.67590 |
| Atu3183 |  |  | -2.71401 |
| Atu0730 |  | Lectin-like protein | -3.47126 |
| Atu2463 |  | DUF306 domain-containing protein | -3.53021 |
| Atu3507 |  | SH3b domain-containing protein | -3.7526 |
| Atu4766 |  | Curlin | -4.14172 |
| Atu3295 |  | Lectin-like protein | -.445499 |
| Atu8200 |  |  | -4.88619 |
| Atu1683 |  |  | -5.14385 |

**Table 2.** Summary of top genes upregulated and downregulated in the Δ*feuP* mutant identified by transcriptome analysis. RNA-seq comparison between *A. tumefaciens* wild-type and the Δ*feuP* mutant revealed genes with an increase or decrease in transcript abundance.

| Locus | Gene Name | Predicted Function | logFC |
| --- | --- | --- | --- |
| Atu1221 |  | NlpC/p60 superfamily protein | 2.74062 |
| Atu1805 |  | BON domain-containing protein | 2.53753 |
| Atu0944 | *cscA* | Sucrose hydrolase | 2.34153 |
| Atu6126 | *traC* | Conjugal transfer protein | 2.22831 |
| Atu2303 |  | Polysaccharide deacetylase | 2.22664 |
| Atu4341 | *impC* | Type VI secretion | 2.04074 |
| Atu4342 | *impB* | Type VI secretion | 2.02102 |
| Atu4339 | *impE* | Type VI secretion | 1.95977 |
| Atu0898 |  | DUF930 domain-containing protein | 1.94824 |
| Atu8089 |  |  | 1.93182 |
| Atu2163 |  | Antifreeze protein | -2.54657 |
| Atu4767 |  | CsgH-like domain-containing protein | -2.86901 |
| Atu0730 |  | Lectin-like protein | -3.22177 |
| Atu4766 |  | Curlin | -3.49563 |
| Atu2463 |  | DUF306 domain-containing protein | -3.63874 |
| Atu1685 |  |  | -3.65474 |
| Atu3507 |  | SH3b domain-containing protein | -3.69132 |
| Atu3295 |  | Lectin-like protein | -4.43674 |
| Atu8200 |  |  | -4.73370 |
| Atu1683 |  |  | -5.09347 |

**Table 3.**
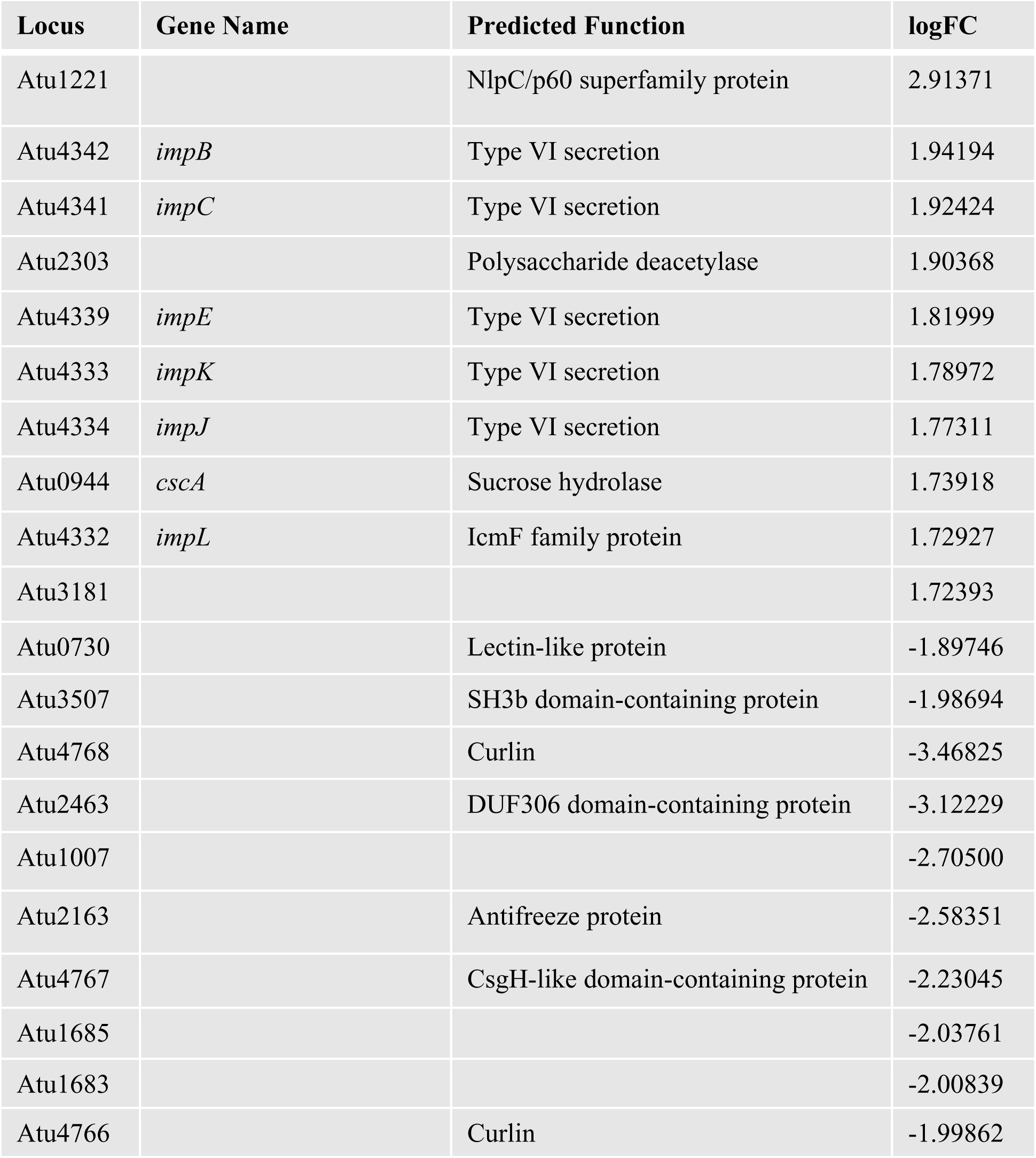
Summary of top genes upregulated and downregulated in the Δ*feuQ* mutant identified by transcriptome analysis. RNA-seq comparison between *A. tumefaciens* wild-type and the Δ*feuQ* mutants revealed genes with an increase or decrease in transcript abundance.

| Locus | Gene Name | Predicted Function | logFC |
| --- | --- | --- | --- |
| Atu1221 |  | NlpC/p60 superfamily protein | 2.91371 |
| Atu4342 | *impB* | Type VI secretion | 1.94194 |
| Atu4341 | *impC* | Type VI secretion | 1.92424 |
| Atu2303 |  | Polysaccharide deacetylase | 1.90368 |
| Atu4339 | *impE* | Type VI secretion | 1.81999 |
| Atu4333 | *impK* | Type VI secretion | 1.78972 |
| Atu4334 | *impJ* | Type VI secretion | 1.77311 |
| Atu0944 | *cscA* | Sucrose hydrolase | 1.73918 |
| Atu4332 | *impL* | IcmF family protein | 1.72927 |
| Atu3181 |  |  | 1.72393 |
| Atu0730 |  | Lectin-like protein | -1.89746 |
| Atu3507 |  | SH3b domain-containing protein | -1.98694 |
| Atu4768 |  | Curlin | -3.46825 |
| Atu2463 |  | DUF306 domain-containing protein | -3.12229 |
| Atu1007 |  |  | -2.70500 |
| Atu2163 |  | Antifreeze protein | -2.58351 |
| Atu4767 |  | CsgH-like domain-containing protein | -2.23045 |
| Atu1685 |  |  | -2.03761 |
| Atu1683 |  |  | -2.00839 |
| Atu4766 |  | Curlin | -1.99862 |

Kyoto Encyclopedia of Genes and Genomes (KEGG) pathway enrichment analysis of differentially expressed genes from each strain background confirmed the large overlap of metabolic pathways and cellular functions regulated by the FeuP-FeuQ system (Fig. 4C, S2). Of the 15 collective pathway classifications generated from the three top 10 pathways for each individual strain background, 36% (5/14) are shared among all three. A total of 71% (10/14) are shared by at least two of the strain backgrounds. Top KEGG pathway classifications shared among all three strain backgrounds include “Metabolic pathways,” “Microbial metabolism in diverse environments,” “ABC transporters,” “Two-component systems,” and “Benzoate degradation.” These results support a role for the FeuP-FeuQ system in regulating bacterial metabolism in addition to more specific functions such as type VI secretion and surface-associated structures referenced above.

### Phenotype profiling supports FeuP-FeuQ regulation of response to osmotic and ionic stress

To further identify physiological roles mediated by the FeuP-FeuQ system, Biolog Phenotype MicroArray (PM) plates were used. PM plates one through twenty (PM1 – PM20), curated by Biolog, are a suite of 96-well plates arrayed with a variety of chemical additions used to assess capacity for growth and metabolism in the presence of these substrates. Growth of the Δ*feuPQ* strain was compared against the wild-type strain after 24 hours, 48 hours, and 72 hours, in duplicate experiments. Growth for both wild-type and Δ*feuPQ* strains was classified for each of 1907 experimental conditions by comparison to that of the wild-type strain at the lowest level of supplementation per experimental condition. Growth that was one doubling less than the control, or lower, was classified as “reduced”. Growth that was one doubling greater than the control, or higher, was classified as “enhanced”. Finally, the growth phenotype (reduced, enhanced, or normal) was compared between the two strains at each time point. Those conditions which gave consistent enhanced or reduced growth, relative to the wild-type growth under the same condition, were identified as potentially regulated by the FeuP-FeuQ system (Table 4).

**Table 4.** Summary of substrate classes and their effect on growth using the Biolog Phenotype MicroArray platform. The number of conditions enhancing or reducing growth of the Δ*feuPQ* strain relative to the wild-type strain is indicated.

| Plate ID | Substrate Class | 24 hours |  | 48 hours |  | 72 hours |  |
| --- | --- | --- | --- | --- | --- | --- | --- |
|  |  | growth enhanced | growth reduced | growth enhanced | growth reduced | growth enhanced | growth reduced |
| PM1 | Carbon utilization | 1/95 | 7/95 | 1/95 | 4/95 | 3/95 | 6/95 |
| PM2 |  | 0/95 | 8/95 | 0/95 | 6/95 | 0/95 | 15/95 |
| PM3 | Nitrogen utilization | 0/95 | 9/95 | 0/95 | 8/95 | 0/95 | 5/95 |
| PM4 | Phosphorous and sulfur utilization | 0/95 | 2/95 | 0/95 | 3/95 | 0/95 | 20/95 |
| PM5 | Biosynthetic pathways/nutrient stimulation | 0/94 | 20/95 | 0/94 | 21/95 | 0/94 | 2/94 |
| PM6 | Nitrogen utilization | 0/94 | 0/94 | 0/94 | 0/94 | 2/94 | 1/94 |
| PM7 |  | 20/94 | 0/94 | 11/94 | 0/94 | 3/94 | 4/94 |
| PM8 |  | 2/94 | 1/94 | 7/94 | 0/94 | 2/94 | 6/94 |
| PM9 | Osmotic/ionic response | 0/96 | 49/96 | 0/96 | 7/96 | 0/96 | 2/96 |
| PM10 | pH response | 4/96 | 0/96 | 19/96 | 0/96 | 4/96 | 0/96 |
| PM11 | Bacterial chemical sensitivity | 3/96 | 5/96 | 2/96 | 3/96 | 3/96 | 5/96 |
| PM12 |  | 3/96 | 6/96 | 3/96 | 1/96 | 0/96 | 7/96 |
| PM13 |  | 2/96 | 2/96 | 6/96 | 1/96 | 0/96 | 6/96 |
| PM14 |  | 13/96 | 0/96 | 2/96 | 5/96 | 0/96 | 16/96 |
| PM15 |  | 1/96 | 9/96 | 1/96 | 9/96 | 2/96 | 26/96 |
| PM16 |  | 6/96 | 6/96 | 1/96 | 11/96 | 1/96 | 29/96 |
| PM17 |  | 5/96 | 5/96 | 1/96 | 8/96 | 0/96 | 12/96 |
| PM18 |  | 4/96 | 6/96 | 3/96 | 4/96 | 0/96 | 22/96 |
| PM19 |  | 1/96 | 16/96 | 5/96 | 3/96 | 1/96 | 33/96 |
| PM20 |  | 2/96 | 15/96 | 3/96 | 4/96 | 2/96 | 6/96 |

A majority of experimental conditions did not differentially affect growth between the wild-type and Δ*feuPQ* strains (Tables 4 and S7). After 24 hours, out of a total of 233 hits (12%), 166 hits showed a reduction in growth, while 67 hits showed an enhancement in growth. After 48 hours, out of 163 total hits (9%), 98 hits resulted in reduced growth, while growth was promoted by 65 hits. After 72 hours, out of a total of 247 hits (27%), 224 hits resulted in a reduction of growth, while only 23 hits resulted in an enhancement of growth. Across all three time points, only 9 hits were shared. Growth was consistently enhanced in the Δ*feuPQ* background in the presence of bleomycin and orphenadrine. Growth was consistently reduced in the Δ*feuPQ* background in the presence of L-phenylalanine, 3-hydroxy-2-butanone, lincomycin, 5,7-dichloro-8-hydroxyquinoline, cetylpyridinium chloride, cinoxacin, and phenylmethylsulfonyl fluoride. The largest set of hits negatively affecting growth of the Δ*feuPQ* strain were found on PM9. This plate contains a set of chemical additions meant to evaluate the response of the organism to osmotic and ionic stresses. Fully 51% (49/96) of experimental conditions on this plate consistently reduced growth after 24 hours. This response was muted at later time points, likely due to longer exposure also inhibiting growth of the wild-type strain. After 72 hours, increased sensitivity of the Δ*feuPQ* strain to chemical inhibitors of metabolism and antimicrobial agents became apparent, with between 5% (5/96) and 34% (33/96) of compounds on plates PM11 – PM20 reducing growth of the mutant strain. The Biolog data matched the broader physiological categories revealed through KEGG enrichment (30, 33, 34).

## DISCUSSION

### FeuP-FeuQ is a conserved regulatory system enabling efficient host interactions

This work establishes the FeuP-FeuQ two-component system as required for efficient surface attachment, motility, and virulence of *A. tumefaciens*. Homologues of FeuP and FeuQ have been studied in several additional members of the *Rhizobiaceae*. In some instances, these homologues are annotated as PhoP (for FeuP) and PopQ (for FeuQ), although synteny and homology support their identification as *bona fide* FeuP or FeuQ homologues (41, 46). FeuP-FeuQ was first recognized as promoting iron acquisition and competitiveness in the rhizosphere in *Rhizobium leguminosarum* (47, 48). Later work established a role for FeuP-FeuQ of *Sinorhizobium meliloti* in the response of this bacterium to osmotic stress. In particular, production of cyclic-β-glucans was demonstrated to be under FeuP-FeuQ regulatory control (41). Furthermore, impaired activity of the FeuP-FeuQ system resulted in reduced nodulation of alfalfa by *S. meliloti*. More recently, transposon-based fitness screening identified the *A. tumefaciens* FeuP-FeuQ homologues as required for efficient survival within tumors formed on tomato (56). Our genetic and phenotypic analyses support the model that *A. tumefaciens* FeuP-FeuQ promotes behaviors required for productive surface engagement, host interaction, and rhizosphere survival.

### Phenotype and transcriptome profiling support FeuP-FeuQ as regulating the response to cell envelope stress

Biolog Phenotype MicroArray plates revealed that loss of FeuP-FeuQ activity results in differential growth and survival kinetics on a wide range of substrates (Table 4), indicating broad physiological impacts of this system (33–36). Particularly notable were increased sensitivities to osmotic, ionic, and antimicrobial stresses. In parallel, RNA-seq identified overlapping and strain-specific DEGs (Fig. 4) regulated by FeuP-FeuQ, further described below. Since a significant number of genes were uniquely altered in only one or two strains, some genes may be regulated independently by *feuP* or only by *feuQ*. The functional categories among DEGs align with known Alphaproteobacterial stress response regulons (28, 34–35). The Δ*feuPQ* mutant displayed the greatest magnitude and number of transcriptional changes, consistent with potential independent and combined regulatory control by FeuP and FeuQ. These results are also consistent with prior metabolic frameworks described for *Alphaproteobacteria* (33, 35–37). The selective transcriptomic changes also raise the possibility of cross-talk between FeuP-FeuQ and additional regulatory pathways (35, 37–38). These may include two-component systems such as ChvG-ChvI or the PdhS-DivK-CtrA pathway, which are known to influence a multitude of phenotypes for survival. Integration of these pathways may enable *A. tumefaciens* to dynamically toggle between behaviors in response to complex environmental cues, especially those encountered during host interaction or niche transitions (17, 28, 38).

### FeuP-FeuQ contributes to host-associated outcomes

Disruption of either *feuP* or *feuQ* altered both biofilm formation and swimming motility without detectable effects on growth (Fig. 2, Fig. S1). The parallel decrease in motility and attachment in the absence of FeuP-FeuQ activity suggests independent regulation of both processes, as opposed to directly regulating the motile-to-sessile transition. This is suggestive of global regulation of cell physiology. Loss of FeuP-FeuQ signaling also reduced tumor formation on potato (Fig. 3). Expression of *S. meliloti ndvA*, involved in cyclic-β-glucan biosynthesis, is activated by the *S. meliloti* FeuP-FeuQ system and required for efficient symbiosis. The *A. tumefaciens ndvA* homologue, *chvA* and the related *chvB* gene are both downregulated in our transcriptomic data. Earlier work on *chvA* (*atu2728*) and *chvB* (*atu2730*) identified production and export of β-1,2-glucans as required for efficient attachment to surfaces and for virulence (49, 50). The primary role for these glucans, however, is osmoadaptation. This role is consistent with our phenotypic and transcriptomic data and further suggests regulatory control over a broad range of phenotypes by the FeuP-FeuQ system.

### FeuP-FeuQ promotes efficient motility and surface attachment

The motility defect as measured in the plate-based assay may be due to one or more factors. In this assay, a reduced growth rate could contribute to smaller swim ring diameters in the mutant backgrounds. More direct effects on flagellar-based swimming motility may also contribute to reduced swim ring diameters. In this case, the plate-based assay does not differentiate among defects in chemotaxis (Che), flagellar motor function (Mot), or flagellar structural issues (Fla). Transcriptomic results in this work support reduced expression of *atu0514*, encoding a methyl-accepting chemotaxis protein, in both the Δ*feuPQ* and Δ*feuP* strain backgrounds. Expression of the flagellar genes *atu0568* and *flgB* (*atu0555*) are also reduced in these two strains. Additional flagellar and motility genes with reduced expression in the Δ*feuP* strain background include *fliF* (*atu0523*), *flgC* (*atu0554*), *atu0559*, *motA* (*atu0560*), *motB* (*atu0569*), *motC* (*atu0570*), *atu0572*, and *flhA* (*atu0581*). As we observed no effect on growth rate when the FeuP-FeuQ system was disrupted, the motility phenotype identified appears most likely to be the result of reduced expression of flagellar- and motility-supporting genes.

Surface attachment and biofilm formation in *A. tumefaciens* is a complex phenotype dependent on a number of known adhesive structures such as the unipolar polysaccharide (UPP) and cellulose. Only one UPP-associated gene, *uppK* (*atu2374*), a gene of unknown function was downregulated in our RNA-seq datasets, for the Δ*feuPQ* and Δ*feuP* strain backgrounds. No cellulose biosynthesis or export genes are differentially expressed. In *Caulobacter crescentus*, a related *Alphaproteobacterium*, surface attachment is mediated primarily through an adhesive holdfast located at the tip of the bacterial stalk. Central to anchoring the holdfast is a holdfast anchoring complex comprised of the HfsABD proteins (51, 52). These proteins share homology with curli assembly and export proteins CsgA, CsgG, and an outer membrane associated surface adhesin, respectively. Although not direct homologues of *hfsA*, *hfsB*, or *hfsD*, three *A. tumefaciens* genes characterized as curlin (*atu4766*, *atu4768*) or a CsgH-like domain-containing protein (*atu4767*), predicted to inhibit curli assembly, are among the most highly downregulated genes in all three RNA-seq datasets. It is possible that Atu4766-Atu4768 functions similarly to HfsABD to mediate attachment or release of the unipolar polysaccharide. Thus, FeuP-FeuQ may positively regulate this function to promote surface interactions. Efficient swimming motility is also required for surface attachment and biofilm formation under static and flow conditions, and it is possible that reduced motility as a result of the observed reduced expression of chemotaxis, flagellar, and motility genes contributes to the attachment phenotype when FeuP-FeuQ activity is compromised (19).

### FeuP-FeuQ regulatory control parallels known responses to cell envelope stress

Among the most highly upregulated genes in all three mutant strains evaluated are numerous components of the *A. tumefaciens* type VI secretion system. This system is used by *A. tumefaciens* during plant host colonization to compete against other rhizosphere-associated bacteria (7, 8). In addition, a large number of genes associated with succinoglycan (SCG) production (53) or export are upregulated in the absence of either *feuP* or *feuQ*, or both. In the Δ*feuPQ* strain background, 11 SCG genes are increased in expression: *exoQ*, *exoF*, *exoY*, *exoP*, *exoN*, *exoO*, *exoM*, *exoA*, *exoL*, *exoK*, and *exoV*. In the Δ*feuP* strain background, 10 SCG genes are upregulated, including all genes in the Δ*feuPQ* set except for *exoL*. In the Δ*feuQ* strain background, five SCG genes were upregulated: *exoQ*, *exoF*, *exoY*, *exoN*, and *exoX*. Taken together, upregulation of type VI secretion system and succinoglycan biosynthesis and export genes, at the same time as decreased expression of motility genes, and the associated phenotypes of reduced motility, reduced surface attachment, and reduced virulence, are remarkably similar to the phenotypes and changes in expression associated with the acid and cell wall stress regulons of *A. tumefaciens* (54, 55).

Similar results have also been observed when the ChvG-ChvI periplasmic regulator, ExoR, is impaired (7). Remarkably, microarray-based transcriptome analysis of the Δ*exoR* strain identified *atu1221* as the most highly upregulated gene, in addition to *atu1131* (*aopB*), as the most highly upregulated annotated gene. *atu1221* is the most highly upregulated gene in all three Δ*feuPQ*, Δ*feuP*, and Δ*feuP*, DEG lists in this work, and *atu1131* is also upregulated in all three. Combined with results from our Phenotype MicroArrays indicating increased sensitivity to osmotic and ionic stresses, this work suggests that the FeuP-FeuQ two-component system is an additional mechanism by which *A. tumefaciens* senses and responds to environmental stresses, particularly conditions affecting cell wall or cell membrane integrity. Together, these results position FeuP-FeuQ as a critical signaling system in *A. tumefaciens*, integrating environmental inputs to coordinate behaviors supporting surface colonization and environmental navigation.

## MATERIALS AND METHODS

### Bacterial strains and growth conditions

Bacterial strains and plasmids used in this study are listed in Tables S1–S3. *A. tumefaciens* str. C58 and derivative strains were routinely cultured in AT minimal medium supplemented with glucose and ammonium sulfate (ATGN) [79 mM KH_2_PO_4_, 15 mM (NH_4_)_2_SO_4_, 600 μM MgSO_4_•7H_2_O, 60 μM CaCl_2_•2H_2_O, 7.1 μM MnSO_4_•H_2_O, 1% glucose, 22 μM Fe_2_SO_4_•7H_2_O], at 28 °C. *E. coli* was routinely cultured in lysogeny broth (LB) at 37 °C. Super optimal broth with catabolite repression (SOC) medium was used for recovery following transformations. Under the appropriate conditions, all growth media were supplemented with carbenicillin (100 μg•mL^-1^), gentamycin (30 μg•mL^-1^ for *E. coli* and 150 μg•mL^-1^ *A. tumefaciens*), or kanamycin (30 μg•mL^-1^ and 150 μg•mL^-1^) when antibiotic supplements were required. Isopropyl-β-D-thiogalactopyranoside (IPTG) was added to a final concentration of 250 μM for induction of gene expression from pSRKGm constructs. Media components were purchased from Fisher Scientific Company, LLC (Hanover Park, IL).

### Construction of deletion and complementation strains

Splicing by overlap extension (SOE) PCR was used to generate deletion constructs targeting *feuP* or *feuQ* singly, or the complete *feuPQ* operon. Approximately 500 base pairs (bp) upstream and 500 bp downstream of the targeted sequence were amplified using primers provided in Table S1. Reactions were performed with Phusion DNA polymerase, using *A. tumefaciens* genomic DNA as the template. The upstream and downstream fragments were fused via overlap extension PCR, with a final amplification using flanking primers to generate the full deletion amplicon. The resulting PCR product was A-tailed with *Taq* DNA polymerase and ligated into the pGEM-T Easy vector. This was then transformed into *E. coli* DH10B for propagation. For the *feuP* and *feuQ* deletion constructs, sequence-verified amplicons were subcloned into suicide vector pNPTS138. The *feuPQ* deletion construct was transferred into pNPTS138 via Gibson assembly. pNPTS138 derivatives were transformed into *E. coli* strain S17-1/λ*pir* for conjugation. Constructs were introduced into *A. tumefaciens* strain C58 via biparental conjugation. Transconjugants were selected by plating on ATGN agar containing 300µg•mL^-1^ kanamycin. Transconjugants were counterselected on AT agar containing 5% sucrose (ATSN). Recombinants were patched onto both ATGN plates 300µg•mL^-1^ kanamycin plates and ATSN plates. Km^R^ Suc^S^ colonies were streak purified and confirmed by PCR.

Directional cloning or Gibson assembly was used to generate plasmid-borne alleles of *feuP*, *feuQ*, or the entire *feuPQ* operon in the broad-host range expression plasmid, pSRKGm. These alleles are all situated such that they are under IPTG-inducible transcriptional control by the *P_lac_* promoter. Site directed mutagenesis of pGEM-T Easy-borne wild-type alleles of *feuP* and *feuQ* was performed using primers USP375/USP376 for the Asp51Ala substitution in FeuP and USP373/USP374 for the His255Ala substitution in FeuQ. Following amplification, reactions were treated with DpnI to selectively degrade methylated parental template and the resulting products were transformed into *E. coli* strain DH10B cells. Plasmids were purified from individual transformants and verified by sequencing. Verified mutant constructs were then sub-cloned via directional cloning into pSRKGm. Molecular biology reagents were purchased from New England BioLabs, Inc. (Ipswich, MA). Oligonucleotides were purchased from Integrated DNA Technologies, Inc. (Coralville, IA).

### Growth curves

Bacterial growth kinetics were assessed by measuring optical density at 600 nm (OD_600_), over a total period of 36 hours. The first 12 hours of measurements were taken every 2 hours, consecutively. Hours 24 and 36 were measured as sole time points. 5 mL overnight cultures (with antibiotics and IPTG as required) were grown in ATGN at 28 °C, with aeration. After approximately 22 hours, 5 mL subcultures were generated at an OD_600_ of 0.05, using fresh ATGN. Subcultures were incubated with continuous shaking and OD_600_ readings were recorded every 2 hours, using a spectrophotometer. Cuvettes containing only ATGN media were used as blanks. Each growth experiment was independently repeated for three biological replicates, consisting of three technical replicates to ensure reproducibility.

### Biofilm formation

Biofilm formation was assessed using a static coverslip assay. Starter cultures were grown overnight in ATGN at 28 °C. The following morning, subcultures were inoculated in fresh ATGN, to an OD_600_ of 0.1. These were then allowed to grow at 28 °C with aeration until mid-exponential phase, ∼OD_600_ = 0.25-0.6. Cultures were diluted to an OD_600_ of 0.05 and 3 mL was added to each of four wells per experimental condition in a 12-well untreated polystyrene tissue culture plate. A sterile 22 mm plastic coverslip was inserted vertically into each well, with approximately half of the coverslip submerged in the well. All plates were incubated in a closed container containing a vented bottle of saturated potassium sulfate. Plates were maintained for 48 hours, at room temperature. Following the termination of the incubation period, coverslips were stained by immersion in 0.1% (w/v) crystal violet for 3-5 minutes. Excess stain was removed by washing with distilled H_2_O. Bound crystal violet was solubilized in 1 mL of 33% (v/v) acetic acid. The absorbance of the resulting solution was measured at 600 nm (A_600_) using a BioTek Synergy H1 microplate reader. The culture density was independently measured by determining the OD_600_ of each sample. Biofilm formation was quantified as the ratio A_600_/OD_600_, normalized to the wild-type strain within each experimental replicate. ATGN was supplemented with 22 µM FeSO_4_, antibiotics, and 250 µM IPTG, as required. Each strain was analyzed in three independent biological replicates, each consisting of three technical replicates.

### Swim motility

Gross motility was evaluated by measuring swim ring diameters following inoculation of ATGN plates containing 0.25 - 0.3% agar. Isolated colonies taken directly from a streaked plate were used to inoculate swim plates, using a sterile toothpick. The plates were then incubated at room temperature, in a closed container containing a vented bottle of saturated potassium sulfate. Swim ring diameters were measured daily for a total period of 7 days. ATGN was supplemented with antibiotics and 250 µM IPTG as required. Each strain was analyzed in three independent biological replicates, each consisting of three technical replicates.

### Potato tumor assay

To test the ability of each strain to stably transform plant tissue, a potato disc tumor assay was performed. Potatoes were initially surface sterilized and then cored using a sterile cork borer. The central tissue of each core was sliced into ten discs, approximately 0.5 cm thick. Discs were placed on a petri dish containing 1.5% water agar solution. Overnight cultures of each strain were sub-cultured to an OD_600_ of 0.05, and 10 μL was used to inoculate each of five discs per strain. The agar plates were sealed with Parafilm and stored uninterrupted at room temperature. Tumors were counted on days 14 and 21.

### Biolog Phenotype MicroArrays

Biolog Phenotypic MicroArray (PM) plates were used to evaluate growth under a variety of physiological conditions. Wild-type *A. tumefaciens* str. C58 and its isogenic derivative, Δ*feuPQ*, were grown overnight in ATGN at 28 °C. The following morning, strains were sub-cultured to exponential phase, OD_600_ = 0.6 - 0.8. Cells were harvested, washed in Biolog inoculating fluid (IF-0), and resuspended in IF-0 to reach 15% transmittance. PM plate inoculation procedures were performed as recommended by the manufacturer, with minor modifications. Biolog redox dye G was excluded from all inoculations. Following inoculation with 100 μL prepared inoculum per well, plates were grown statically at 30 °C for 72 hours. OD_600_ was measured at 24, 48, and 72 hours, using a BioTek Synergy H1 microplate reader.

### RNA Sequence analysis of mutant strains

For RNA isolation and sequencing, overnight cultures were incubated at 28 °C, with shaking. The following day, cultures were sub-cultured to an OD600 of 0.05. Cultures were then placed back into the incubator to grow until the OD_600_ reached 0.8 for all strains. Cells were harvested and RNA stabilized by combining 650 μL of culture with 1350 μl of Qiagen RNAprotect reagent. Samples were stored at −20 °C until processed. Total RNA was isolated using the Qiagen RNeasy Mini Kit, per manufacturer’s protocol. Samples were treated with RNase-free Dnase I. The sample purity and concentration were evaluated using a Nandrop and Qubit. Samples were sent to Novogene for sequencing. RNA sample quality control, prokaryotic RNA library preparation, and library sequencing using Illumina paired-end sequencing on a NovaSeq X Plus. Approximately 20 million reads per sample were obtained. FASTQC was used for quality control of reads, followed by trimming with Trimmomatic, and then aligned to the *A. tumefaciens* strain C58 reference genome using Spliced Transcripts Alignment to a Reference (STAR). Differential expression analysis was completed using edgeR.

RNA-seq data have been deposited with the Gene Expression Omnibus under the accession GSE316521 (private until publication).

### Statistical Analysis

In each experiment, all assays were performed, at minimum, in triplicate data sets for each sample (technical replicates). All experiments were repeated at minimum a total of three times (biological replicates). The exception to this procedure is shown in the Biolog phenotype assay, in which the experiment was repeated for a total of two biological replications. Statistical significance was determined using one-way analysis of variance (ANOVA) or Student’s *t*-test, as appropriate.

## ACKNOWLEDGEMENTS

This work was funded by NSF CAREER Award 2238568 and NIFA CAREER Award 2023-67014-40246.

---

**Table S1:** List of Strains used in this study.

| *E. coli* | Strain | Characteristics | Source |
| --- | --- | --- | --- |
| | DH5α/λ*pir* | *endA1 hsdR17 glnV44 (= supE44) thi-1 recA1 gyrA96 relA1* ϕ80*dlac*Δ(*lacZ*)M15 $\Delta$(*lacZYA-argF*)U169 *zdg-232::Tn10 uidA::pir+* | (1) |
| | DH10B | F$-$ *mcrA* $\Delta$(*mrr-hsdRMS-mcrBC*) ϕ80*lacZ*ΔM15 Δ*lacX74 recA1 endA1 araD139* $\Delta$(*ara-leu*)7697 *galU galK* λ-*rpsL(Str<sup>R</sup>) nupG* | (2) |
| | JM109 | *endA1 recA1 gyrA96 thi hsdR17 (r<sub>k</sub>$^-$ m<sub>k</sub>$^+$) relA1 supE44* $\Delta$(*lac-proAB*) [F' *traD36 proAB laqI<sup>q</sup>Z*ΔM15] | (3) |
| *A. tumefaciens* | C58 | Nopaline type strain, pAtC58+, pTiC58+ | (4) |
| | C58-RWN001 | C58 Δ*feuP* (Δ*atu0970*) | This Study |
| | C58-RWN002 | C58 Δ*feuQ* (Δ*atu0971*) | This Study |
| | C58-RWN011 | C58 Δ*feuPQ* (Δ*atu0970* Δ*atu0971*) | This Study |

**Table S2:**
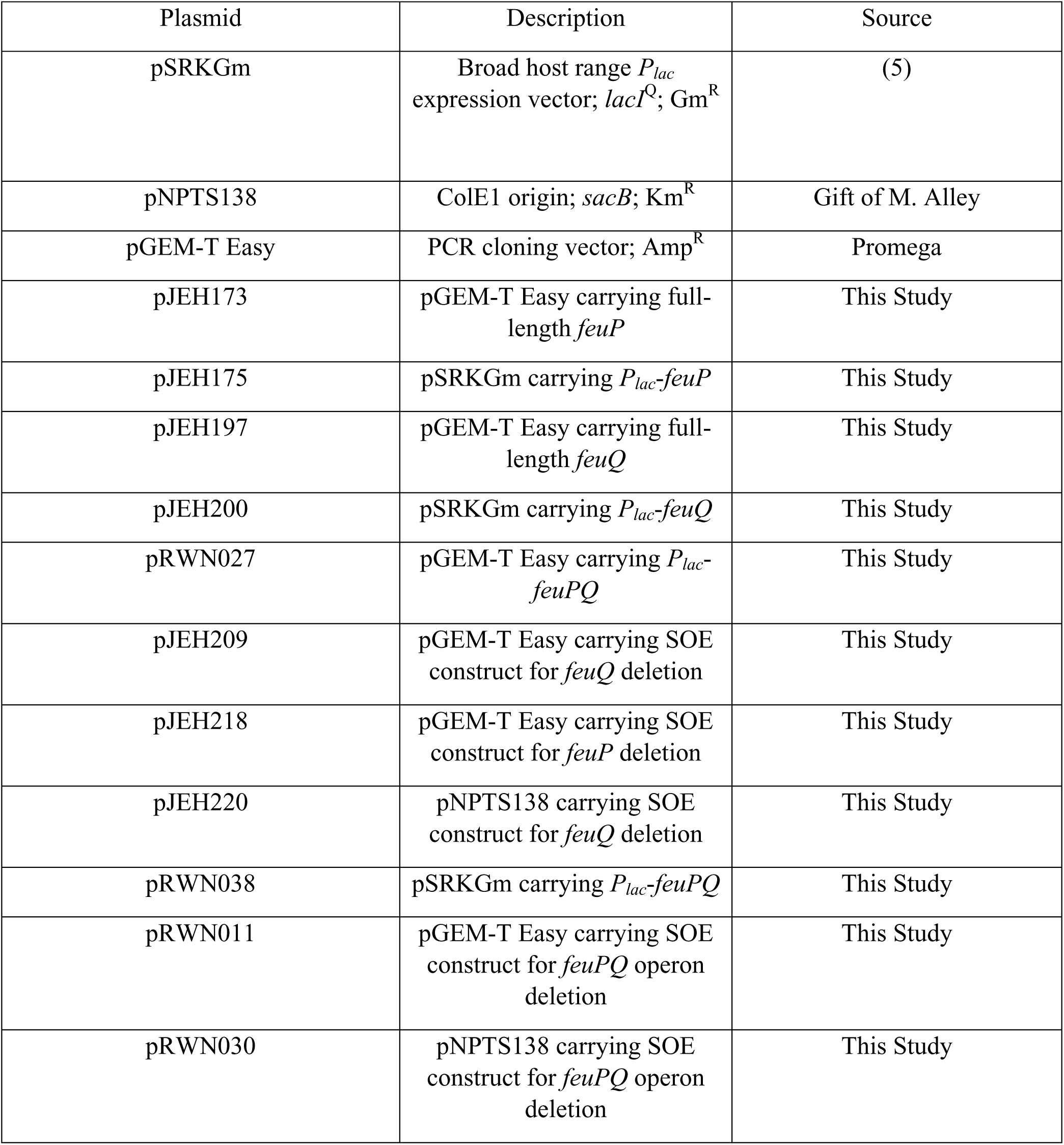
List of plasmids in this study.

**Table S3:**
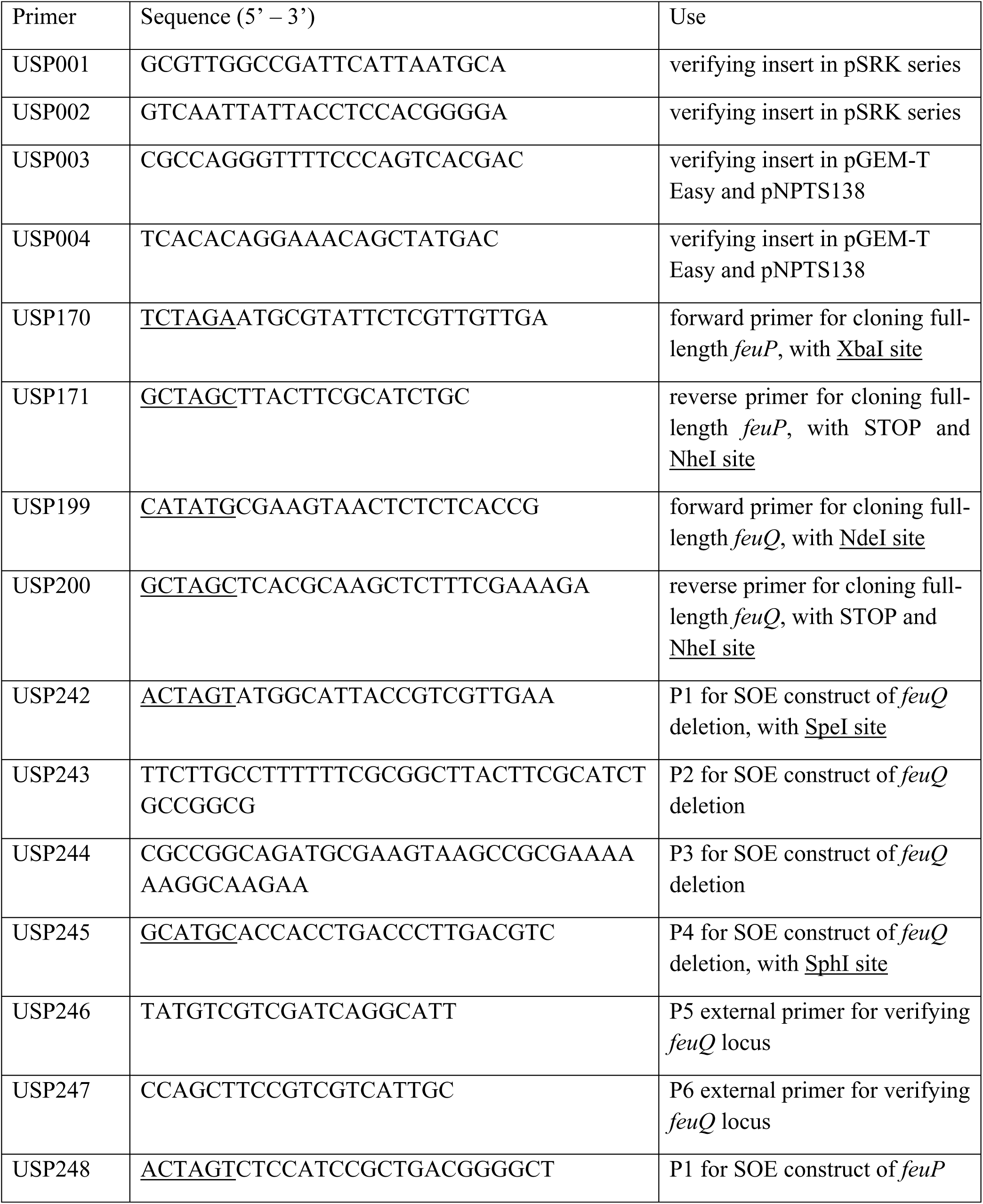
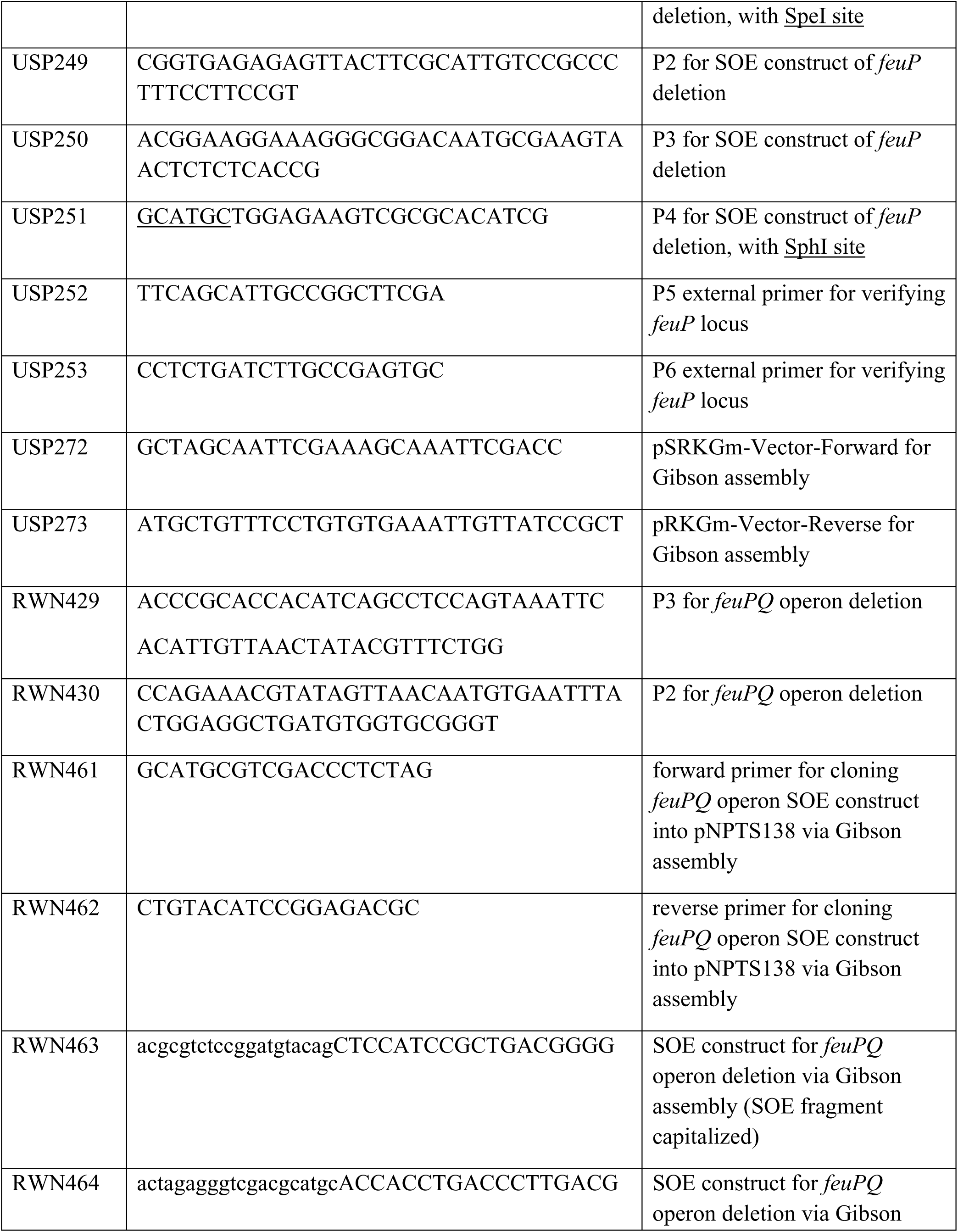
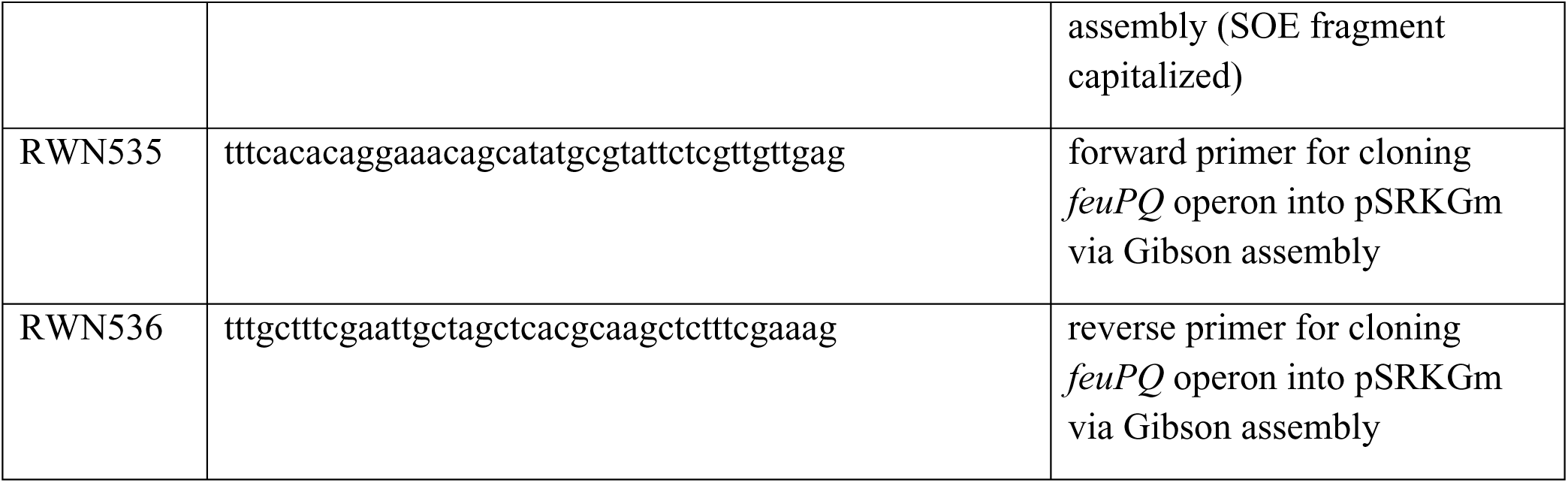
List of DNA primers used in this study.

**Fig S1.**
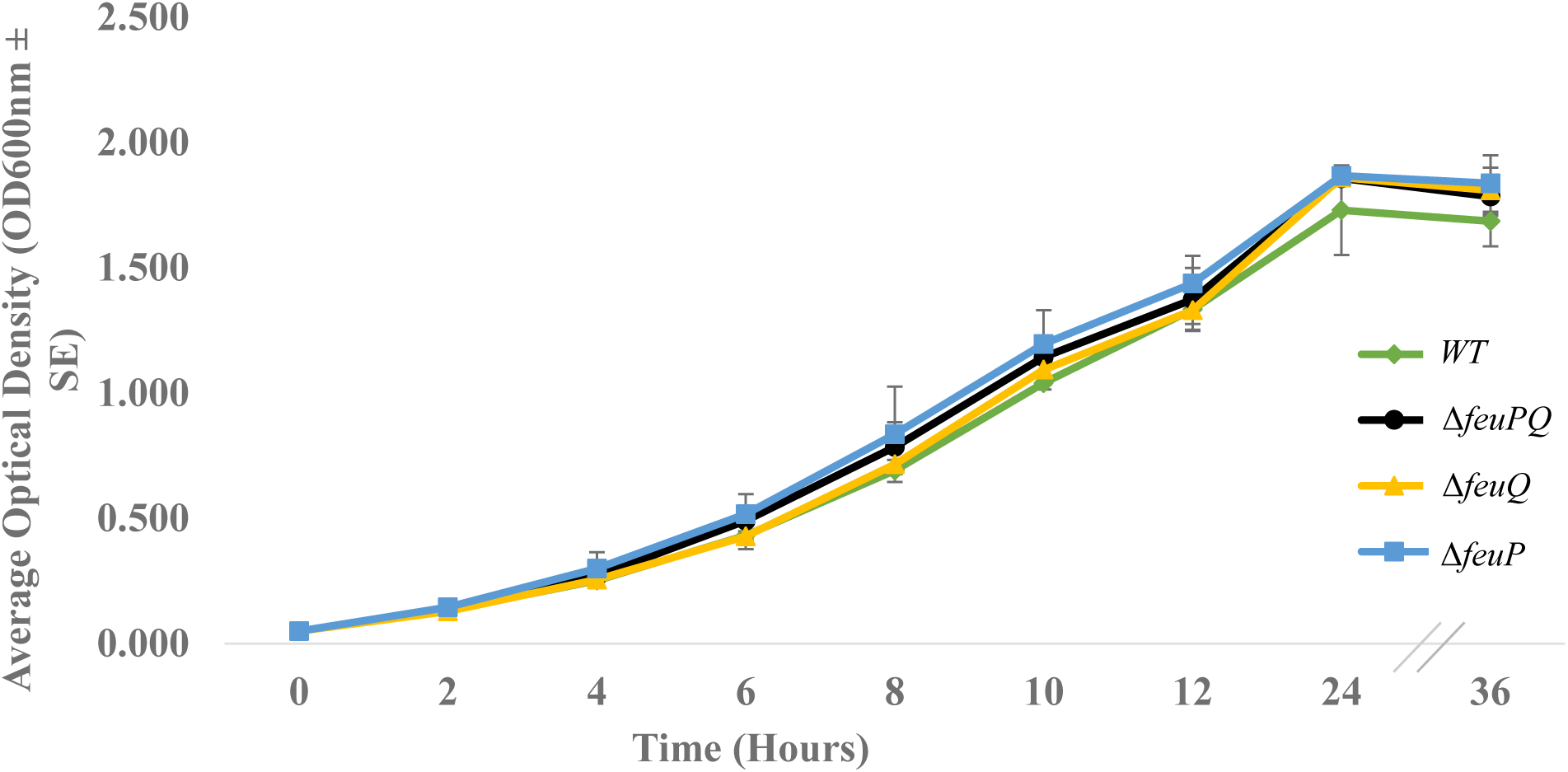
**Loss of *feuP* or *feuQ*, singly or in combination, does not significantly affect growth rate.** Growth of wild-type *A. tumefaciens* strain C58 (WT) and the indicated derivatives in ATGN medium was monitored by OD_600,_ over a total period of 36 hours. Data shown are means ± standard error from three independent experiments, each of which contained three biological replicates. No statistically significant differences were detected among strains by one-way ANOVA, where p< 0.05.

**Fig S2.**
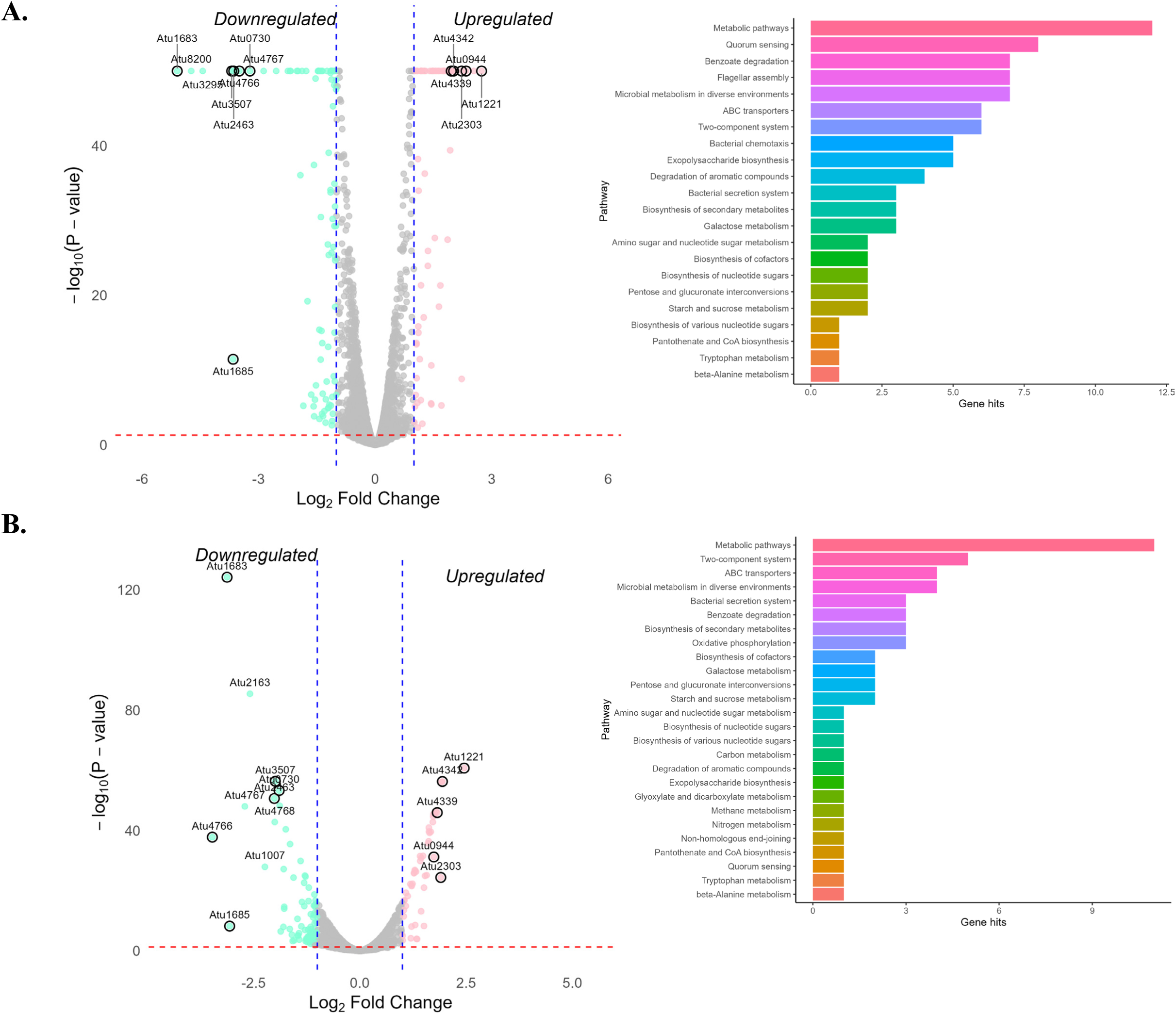
Volcano plots and KEGG pathway assignments for differentially expressed genes in Δ*feuP* and Δ*feuQ* derivatives of *A. tumefaciens*. Volcano plots, left, and KEGG pathway assignments, right, show the suite of genes differentially expressed, relative to the wild-type strain, for the Δ*feuP* (A) and Δ*feuQ* (B) strains.

**Fig S3.**
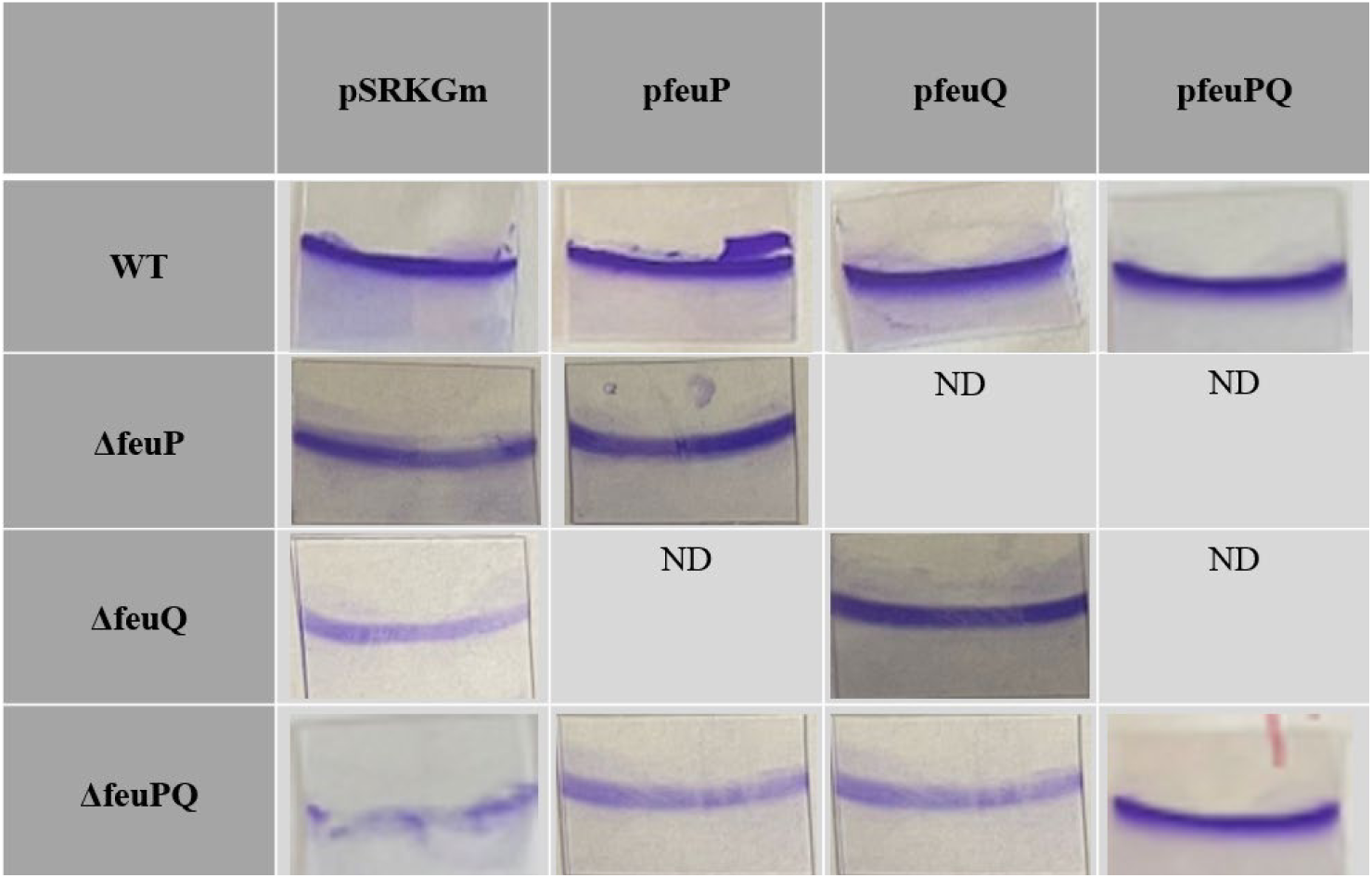
**Loss of *feuP* or *feuQ*, singly or in combination, significantly alters surface attachment**. Crystal violet stained coverslips from the static biofilm assay quantified in Figure 2 are shown. ND, not done.

**Fig S4.**
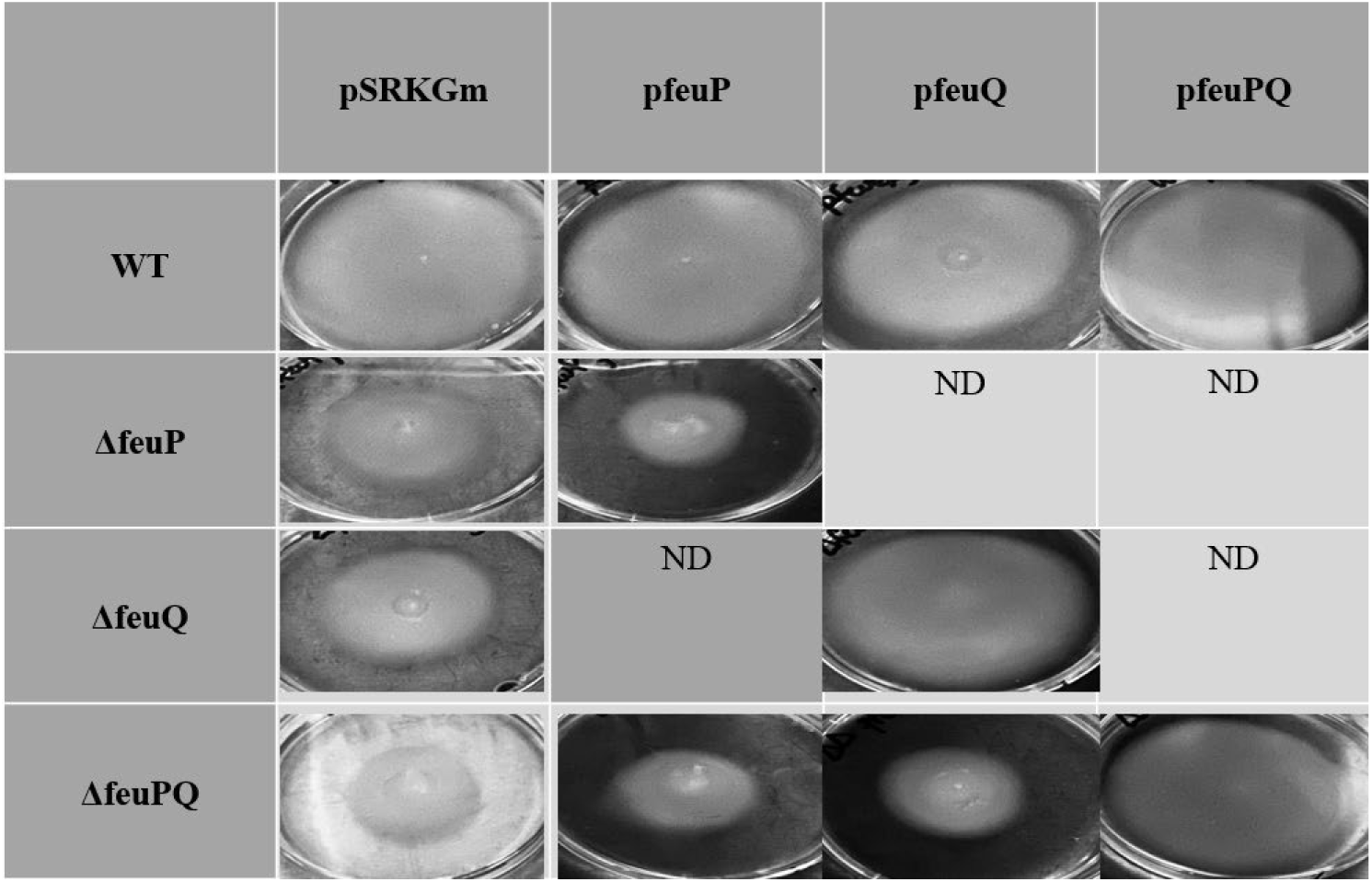
Loss of *feuP* or *feuQ*, singly or in combination, significantly alters swim motility. Swim plates on day 7 from the motility assay quantified in Figure 2 are shown. ND, not done.

## References

1. Stock AM, Robinson VL, Goudreau PN. 2000. Two-component signal transduction. Annu Rev Biochem 69: 183–215.

2. Skerker JM, Prasol, MS, Perchuk BS, Biondi EG, Laub MT. 2005. Two-component signal transduction pathways regulating growth and cell cycle progression in a bacterium: a system-level analysis. PLoS Biol 3(10): e334.

3. Matthysse AG. 2014. Attachment of Agrobacterium to plant surfaces. Front Plant Sci 5: 252.

4. Xu J, Kim J, Koestler BJ, Choi JH, Waters CM, Fuqua C. 2013. Genetic analysis of Agrobacterium tumefaciens unipolar polysaccharide production reveals complex integrated control of the motile-to-sessile switch. Mol Microbiol 89(5): 929–948.

5. Thompson MA, Onyeziri MC, Fuqua C. 2018. Function and Regulation of Agrobacterium tumefaciens Cell Surface Structures that Promote Attachment. Curr Top Microbiol Immunol 418: 143–184.

6. Wise AA, Fang F, Lin YH, He F, Lynn DG, Binns AN. 2010. The receiver domain of hybrid histidine kinase VirA: an enhancing factor for vir gene expression in Agrobacterium tumefaciens. J Bacteriol 192(6): 1534–1542.

7. Heckel BC, Tomlinson AD, Morton ER, Choi JH, Fuqua C. 2014. Agrobacterium tumefaciens exoR controls acid response genes and impacts exopolysaccharide synthesis, horizontal gene transfer, and virulence gene expression. J Bacteriol 196(18): 3221–3233.

8. Wu CF, Lin JS, Shaw GC, Lai EM. 2012. Acid-induced type VI secretion system is regulated by ExoR-ChvG/ChvI signaling cascade in Agrobacterium tumefaciens. PLoS Pathog 8(9): e1002938.

9. Alakavuklar MA, Heckel BC, Stoner AM, Stembel JA, Fuqua C. 2021. Motility control through an anti-activation mechanism in Agrobacterium tumefaciens. Mol Microbiol 116(5): 1281–1297.

10. Kim J, Heindl JE, Fuqua C. 2013. Coordination of division and development influences complex multicellular behavior in *Agrobacterium tumefaciens*. PLoS one 8(2): e56682.

11. Heindl, JE, Crosby D, Brar S, Pinto JF, Singletary T, Merenich D, Eagan JL, Buechlein AM, Bruger EL, Waters CM, Fuqua C. 2019. Reciprocal control of motility and biofilm formation by the PdhS2 two-component sensor kinase of Agrobacterium tumefaciens. Microbiology (Reading) 165(2): 146–162.

12. Bochner BR. 2009. Global phenotypic characterization of bacteria. FEMS Microbiol Rev 33(1): 191–205.

13. Bourret RB. 2010. Receiver domain structure and function in response regulator proteins. Curr Opin Microbiol 13(2): 142–149.

14. Greenwich JL, Heckel BC, Alakavuklar MA, Fuqua C. 2003. The ChvG–ChvI regulatory network: A conserved global regulatory circuit among the Alphaproteobacterial with pervasive impacts on host interactions and diverse cellular processes. Annual Review of Microbiology 77:131–148.

15. Marina A, Waldburger CD, Hendrickson WA 2005. Structure of the entire cytoplasmic portion of a sensor histidine-kinase protein. EMBO J 24(24): 4247–4259.

16. Gelvin SB. 2012. Traversing the cell: *Agrobacterium* T-DNA’s journey to the host genome. Frontiers in plant science 3(52).

17. Matthysse AG, Marry M, Krall L, Kaye M, Ramey BE, Fuqua C, White AR. 2005. The effect of cellulose overproduction on binding and biofilm formation on roots by Agrobacterium tumefaciens. Mol Plant Microbe Interact 18(9): 1002–1010.

18. Matthysse AG, Thomas DL, White AR. 1995. Mechanism of cellulose synthesis in Agrobacterium tumefaciens. J Bacteriol 177(4): 1076–1081.

19. Merritt PM, Danhorn T, Fuqua C. 2007. Motility and chemotaxis in Agrobacterium tumefaciens surface attachment and biofilm formation. J Bacteriol 189(22): 8005–8014.

20. Li L, Jia Y, Hou Q, Charles TC, Nester EW, Pan SQ. 2002. A global pH sensor: Agrobacterium sensor protein ChvG regulates acid-inducible genes on its two chromosomes and Ti plasmid. Proc Natl Acad Sci U S A 99(19): 12369–12374.

21. Lin JS, Wu HH, Hsu PH, Ma LS, Pang YY, Tsai MD, Lai EM. 2014. Fha interaction with phosphothreonine of TssL activates type VI secretion in Agrobacterium tumefaciens. PLoS Pathog 10(3): e1003991.

22. Ma LS, Narberhaus F, Lai EM. 2012. IcmF family protein TssM exhibits ATPase activity and energizes type VI secretion. J Biol Chem 287(19): 15610–15621.

23. Tomlinson AD, Fuqua C. 2009. Mechanisms and regulation of polar surface attachment in Agrobacterium tumefaciens. Curr Opin Microbiol 12(6): 708–714.

24. Tomlinson AD, Ramey-Hartung B, Day TW, Merritt PM, Fuqua C. 2010. Agrobacterium tumefaciens ExoR represses succinoglycan biosynthesis and is required for biofilm formation and motility. Microbiology (Reading) 156(Pt 9): 2670–2681.

25. Wuichet K, Cantwell BJ, Zhulin IB. 2010. Evolution and phyletic distribution of two-component signal transduction systems. Curr Opin Microbiol 13(2): 219–225.

26. Williams KP, Sobral BW, Dickerman AW. 2007. A robust species tree for the alphaproteobacteria. J Bacteriol 189(13): 4578–4586.

27. Hordt A, Lopez MG, Meier-Kolthoff JP, Schleuning M, Weinhold LM, Tindall BJ, Gronow S, Kyrpides NC, Woyke T, Goker M. 2020. Analysis of 1,000+ Type-Strain Genomes Substantially Improves Taxonomic Classification of Alphaproteobacteria. Front Microbiol 11: 468.

28. Mitrophanov AY, Groisman EA. 2008. Signal integration in bacterial two-component regulatory systems. Genes Dev 22(19): 2601–2611.

29. Kinoshita-Kikuta E, Kusamoto H, Ono S, Akayama K, Eguchi Y, Igarashi M, Okajima T, Utsumi R, Kinoshita E, Koike T. 2019. Quantitative monitoring of His and Asp phosphorylation in a bacterial signaling system by using Phos-tag Magenta/Cyan fluorescent dyes. Electrophoresis 40(22): 3005–3013.

30. van Opijnen T, Camilli A. 2013. Transposon insertion sequencing: a new tool for systems-level analysis of microorganisms. Nat Rev Microbiol 11(7): 435–442.

31. Wadhwa N, Berg HC. 2022. Bacterial motility: machinery and mechanisms. Nat Rev Microbiol 20(3): 161–173.

32. Xu J, Kim J, Danhorn T, Merritt PM, Fuqua C. 2012. Phosphorus limitation increases attachment in Agrobacterium tumefaciens and reveals a conditional functional redundancy in adhesin biosynthesis. Res Microbiol 163(9-10): 674–684.

33. Imam S, Noguera DR, Donohue TJ. 2015. CceR and AkgR regulate central carbon and energy metabolism in alphaproteobacteria. mBio 6(1).

34. Heindl JE, Hibbing ME, Xu J, Natarajan R, Buechlein AM, Fuqua C. 2015. Discrete Responses to Limitation for Iron and Manganese in Agrobacterium tumefaciens: Influence on Attachment and Biofilm Formation. J Bacteriol 198(5): 816–829.

35. Heindl JE, Wang Y, Heckel BC, Mohari B, Feirer N, Fuqua C. 2014. Mechanisms and regulation of surface interactions and biofilm formation in Agrobacterium. Front Plant Sci 5: 176.

36. Imhoff JF. 2006. Phototrophic alpha-proteobacteria. In The prokaryotes. Springer 41–64.

37. Winans SC. 1992. Two-way chemical signaling in Agrobacterium-plant interactions. Microbiol Rev 56(1): 12–31.

38. Chang CH, Winans SC. 1992. Functional roles assigned to the periplasmic, linker, and receiver domains of the Agrobacterium tumefaciens VirA protein. J Bacteriol 174(21): 7033–7039.

39. Brencic A, Winans SC. 2005. Detection of and response to signals involved in host-microbe interactions by plant-associated bacteria. Microbiol Mol Biol Rev 69(1): 155–194.

40. Wu CF, Lin JS, Shaw GC, Lai EM. 2012. Acid-induced type VI secretion system is regulated by ExoR-ChvG/ChvI signaling cascade in Agrobacterium tumefaciens. PLoS Pathog 8(9): e1002938

41. Griffitts JS, Carlyon RE, Erickson JH, Moulton JL, Barnett MJ, Toman CJ, Long SR. 2008. A *Sinorhizobium meliloti* osmosensory two-component system required for cyclic glucan export and symbiosis. Molecular Microbiology 69(5):1230–1246

42. Waldburger L, Thompson MG, Weisberg AJ, Lee N, Chang JH, Keasling JD, Shih PM. 2023. Transcriptome architecture of the three main lineages of agrobacteria. Msystems 8(4), e00333–23.

43. Gao R, Stock AM. 2009. Biological insights from structures of two-component proteins. Annu Rev Microbiol 63: 133–154

44. Kabbara S, Hérivaux A, Dugé de Bernonville T, Courdavault V, Clastre M, Gastebois A, Osman M, Hamze M, Cock JM, Schaap P, Papon N. 2019. Diversity and evolution of sensor histidine kinases in eukaryotes. Genome biology and evolution 11(1):86–108

45. Curtis PD, Bru, YV. 2014. Identification of essential alphaproteobacterial genes reveals operational variability in conserved developmental and cell cycle systems. Molecular microbiology. 93(4):713–735

46. Karp, PD, Billington R, Caspi R, Fulcher CA, Latendresse M, Kothari A, Keseler IM, Midford PE, Ong Q, Ong WK, Paley SM and Subhraveti, P. 2019. The BioCyc collection of microbial genomes and metabolic pathways. Briefings in bioinformatics 20(2019):1085–1093.

47. Yeoman KH, Delgado MJ, Wexler M, Downie JA, Johnston AW. 1997. High affinity iron acquisition in Rhizobium leguminosarum requires the cycHJKL operon and the feuPQ gene products, which belong to the family of two-component transcriptional regulators. Microbiology 143(1):127–134

48. Knights HE, Ramachandran VK, Jorrin B, Ledermann R, Parsons JD, Aroney ST, Poole PS. 2024. Rhizobium determinants of rhizosphere persistence and root colonization. The ISME Journal 18(1):wrae072

49. Xu J, Kim J, Danhorn T, Merritt PM, Fuqua C. 2012. Phosphorus limitation increases attachment in Agrobacterium tumefaciens and reveals a conditional functional redundancy in adhesin biosynthesis. Research in microbiology. 163(9-10): 674–684

50. Cangelosi GA, Martinetti G, Leigh JA, Lee CC, Theines C, Neste EW. 1989. Role for [corrected] Agrobacterium tumefaciens ChvA protein in export of beta-1, 2-glucan. Journal of bacteriology. 171(3):1609–1615

51. Sulkowski NI, Hardy GG, Brun YV, Bharat TA. 2019. A multiprotein complex anchors adhesive holdfast at the outer membrane of Caulobacter crescentus. Journal of Bacteriology. 201(18), 10–1128.

52. Hardy GG, Allen RC, Toh E, Long M, Brown PJ, Cole-Tobian JL, Brun YV. 2010. A localized multimeric anchor attaches the Caulobacter holdfast to the cell pole. Molecular microbiology. 76(2):409–427.

53. Wu D, Li A, Ma F, Yang J, Xie Y. 2016. Genetic control and regulatory mechanisms of succinoglycan and curdlan biosynthesis in genus Agrobacterium. Applied microbiology and biotechnology. 100(14):6183–6192.

54. Williams MA, Bouchier JM, Mason AK, Brown PJ. 2022. Activation of ChvG-ChvI regulon by cell wall stress confers resistance to β-lactam antibiotics and initiates surface spreading in Agrobacterium tumefaciens. PLoS Genetics. 18(12):e1010274.

55. Yuan ZC, Liu P, Saenkham P, Kerr K, Nester EW. 2008. Transcriptome profiling and functional analysis of Agrobacterium tumefaciens reveals a general conserved response to acidic conditions (pH 5.5) and a complex acid-mediated signaling involved in Agrobacterium-plant interactions. Journal of Bacteriology. 190(2):494–507.

56. Torres M, Jiquel A, Jeanne E, Naquin D, Dessaux Y, Faure D. 2022. Agrobacterium tumefaciens fitness genes involved in the colonization of plant tumors and roots. New Phytologist. 233(2): 905–918.

## References

1. Platt, R., C. Drescher, S. K. Park and G. J. Phillips (2000). “Genetic system for reversible integration of DNA constructs and lacZ gene fusions into the Escherichia coli chromosome.” Plasmid 43(1): 12–23.

2. Grant, S. G., J. Jessee, F. R. Bloom and D. Hanahan (1990). “Differential plasmid rescue from transgenic mouse DNAs into Escherichia coli methylation-restriction mutants.” Proc Natl Acad Sci U S A 87(12): 4645–4649.

3. Yanisch-Perron, C., J. Vieira and J. Messing (1985). “Improved M13 phage cloning vectors and host strains: nucleotide sequences of the M13mp18 and pUC19 vectors.” Gene 33(1): 103–119.

4. Watson, B., T. C. Currier, M. P. Gordon, M. D. Chilton and E. W. Nester (1975). “Plasmid required for virulence of Agrobacterium tumefaciens.” J Bacteriol 123(1): 255–264.

5. Khan, S. R., J. Gaines, R. M. Roop, 2nd and S. K. Farrand (2008). “Broad-host-range expression vectors with tightly regulated promoters and their use to examine the influence of TraR and TraM expression on Ti plasmid quorum sensing.” Appl Environ Microbiol 74(16): 5053–5062.

